# Fetal liver neutrophils are responsible for the postnatal neutrophil surge

**DOI:** 10.1101/2024.08.26.609612

**Authors:** Ryo Ishiwata, Yuji Morimoto

**Author notes:** Corresponding Author: Ryo Ishiwata, Department of Physiology, National Defense Medical College, 3-2, Namiki, Tokorozawa City, Saitama Pref., Japan, 359-8513, Tel; (81)4-2995-1211.

## Abstract

Mammalian neonates experience an abrupt surge of blood neutrophil count within the first day of life. The postnatal neutrophil surge is regarded as a defensive reaction against infection; however, the mechanisms underlying this surge remain unclear.

The present study demonstrates that the postnatal neutrophil surge arises from the liver neutrophil pool. In rat neonates, the neutrophil surge was evident at 6 hours after birth. The proportion and immaturity of bone marrow neutrophils remained unaltered at 6 hours but increased only after the surge had peaked. In the rat fetal and neonatal livers, we observed prenatal neutrophil accumulation and acute loss of the neutrophils coinciding with the postnatal neutrophil surge. In *Lys-EGFP* mice, an acute loss of liver neutrophils was observed within 12 hours of birth. This loss was characterized by a decrease in mature neutrophils and by perivascular neutrophil localization in the livers. Additionally, mouse fetuses exhibited an accumulation of the liver neutrophil pool during the late gestational period (e15-18), which was attributable to neutrophil-biased myeloid differentiation mediated by granulocyte-colony stimulating factor (G-CSF). The liver neutrophils exhibited characteristic transcriptomic alterations within three hours of birth, exemplified by an increase in the *Nos*2 (iNOS) gene. The administration of a non-selective NOS inhibitor or an iNOS-selective inhibitor resulted in the inhibition of the postnatal neutrophil surge in rat neonates, accompanied by the retention of liver neutrophils.

These findings shed light on the previously unidentified source of the postnatal neutrophil surge and the stimulus initiating it.

**Significance statement:** Infections in newborns, particularly those occurring within the first 72 hours of life, are leading cause of mortality and morbidity. Neutrophil, a type of leukocytes, acutely increases within 24 hours after birth in the newborns’ blood. This neutrophil surge is regarded as an innate defensive system against infection; however, the mechanisms of the surge have remained unknown. Here, we examined rats and mice and found that the neutrophils accumulated in the fetal livers during the late pregnancy and were released into blood after birth. We also found a specific factor causing the release of the liver neutrophils. These findings might explain why preterm or low-birth weight newborns often lack the postnatal neutrophil surge and are thus more susceptible to infections.

## Introduction

At the time of birth, neonates are required to adapt to the rapid transition from the sterile intraamniotic environment to the external world. Neutrophils constitute the initial line of defense against infection. During the first day of life, neonates undergo a significant increase in circulating blood neutrophils, known as the postnatal neutrophil surge. In human neonates, the blood neutrophil count is 2,000-6,000/μL at birth, which then increases to 7,800-14,500/μL within six to 12 hours after birth (1). Subsequently, the neutrophil count declines gradually over the 72 hours following birth, reaching a value comparable to that of an adult (1). The postnatal neutrophil surge has also been documented in mice (2), indicating a conserved mammalian response.

A deeper comprehension of the neutrophil dynamics at the time of birth may prove instrumental in the prevention of neonatal infection. Neonatal sepsis, particularly those cases occurring within the first 72 hours of life (early-onset sepsis, or EOS), represents a significant cause of morbidity and mortality (3). Additionally, it is associated with the absence of the postnatal neutrophil surge, or neutropenia. (4). In the majority of cases of EOS, neutropenia preceded the onset of sepsis (4), indicating that the neutropenia is a predisposing factor for neonatal sepsis. The reported risk factors for EOS (5), including very-low-birth-weight, caesarean delivery, and male sex, are associated with an absence or lower level of the postnatal neutrophil surge (6, 7). Notably, only 16% of the very-low-birth-weight infants exhibited a proper level of the postnatal neutrophil surge (7).

The available evidence suggests that the postnatal neutrophil surge represents an innate defensive mechanism of mammalian neonates. Nevertheless, the precise mechanisms underlying the postnatal neutrophil surge remain unclear. In instances of adult systemic infection, bone marrow granulopoiesis is selectively activated, resulting in a rapid surge in circulating neutrophils (emergency granulopoiesis). The bone marrow of neonates contains a limited number of proliferative neutrophil pool relative to their body weight. In full-term rat neonates, the neutrophil pool represents only 10% of the value observed in adults (8). Furthermore, an increase in the proportion of immature neutrophils within circulating neutrophils, a feature observed in the emergency granulopoiesis (9), was not observed in the human postnatal neutrophil surge (1). Consequently, the postnatal neutrophil surge is postulated to originate from the extramedullary neutrophil pools, although the precise source remains unknown (10).

The present study demonstrates that the postnatal neutrophil surge originates from the liver rather than the bone marrow. The postnatal neutrophil surge was also observed in rat neonates, with the earliest evidence appearing at 6 hours after birth and the peak occurring at 10 hours. This pattern resembles that observed in human neonates. The proportion of bone marrow neutrophils remained unaltered at the six-hour postnatal time point. In contrast, we observed a prenatal accumulation of liver neutrophils, which was rapidly followed by a decrease after birth. The analysis of *Lys-EGFP* and wild-type mouse fetuses and neonates provided evidence supporting the liver origin of the postnatal neutrophil surge and indicated that the neutrophil-biased myeloid differentiation is promoted prenatally in a G-CSF-dependent manner. In the postnatal period, the liver neutrophil underwent changes in transcriptomic profile within three hours after birth, characterized by an increase in the *Nos2* (iNOS) gene. The administration of both a non-selective NOS inhibitor and an iNOS-selective inhibitor resulted in the inhibition of the postnatal neutrophil surge in rats, accompanied by a retention of liver neutrophils. These results demonstrate that the postnatal neutrophil surge is composed of the prenatal accumulation of liver neutrophils and the postnatal release of liver neutrophils, which are mediated by iNOS.

## Results

### The postnatal neutrophil surge was observed in rat neonates

The relatively small size of mouse neonates represents a significant limitation in the study of fetal and neonatal blood. Therefore, the initial analysis was conducted on rat fetuses and neonates. Figure S1A and B illustrate the typical flow cytometry plots for the blood of a full-term rat fetus (embryonic day 21, e21) and an adult rat (the dam of the fetus). The fetal blood similarly exhibited a typical leukocyte plot pattern of forward scatter (FSC) and side scatter (SSC). Two anti-rat granulocyte antibodies, RP-1 and HIS48, were employed for the detection of neutrophils. Within the CD45^+^ leukocyte population, RP-1 exhibited a specific labeling of SSC^hi^ neutrophils, while HIS48 demonstrated labeling of both neutrophils and monocytes. Fluorescence minus one (FMO) control analyses were conducted to confirm the specificity of each fluorescence signal for each antibody (Fig. S2A and B). Cell sorting and Wright-Giemsa staining were performed to confirm the identity of the cells. The results demonstrated that CD45^+^/RP-1^+^ cells were neutrophils, CD45^+^/RP-1^−^/HIS48^+^ cells were monocytes, and CD45^+^/RP-1^−^/HIS48^−^ cells were lymphocytes (Fig. 1A). The HIS48^hi^ monocytes and the HIS48^int^ monocytes are considered to be classical and non-classical monocytes, respectively (11). The fetal blood sample contained only a small population of HIS48^int^ monocytes, which was distinctive from the adult blood sample (Fig. S1).

**Figure 1.**
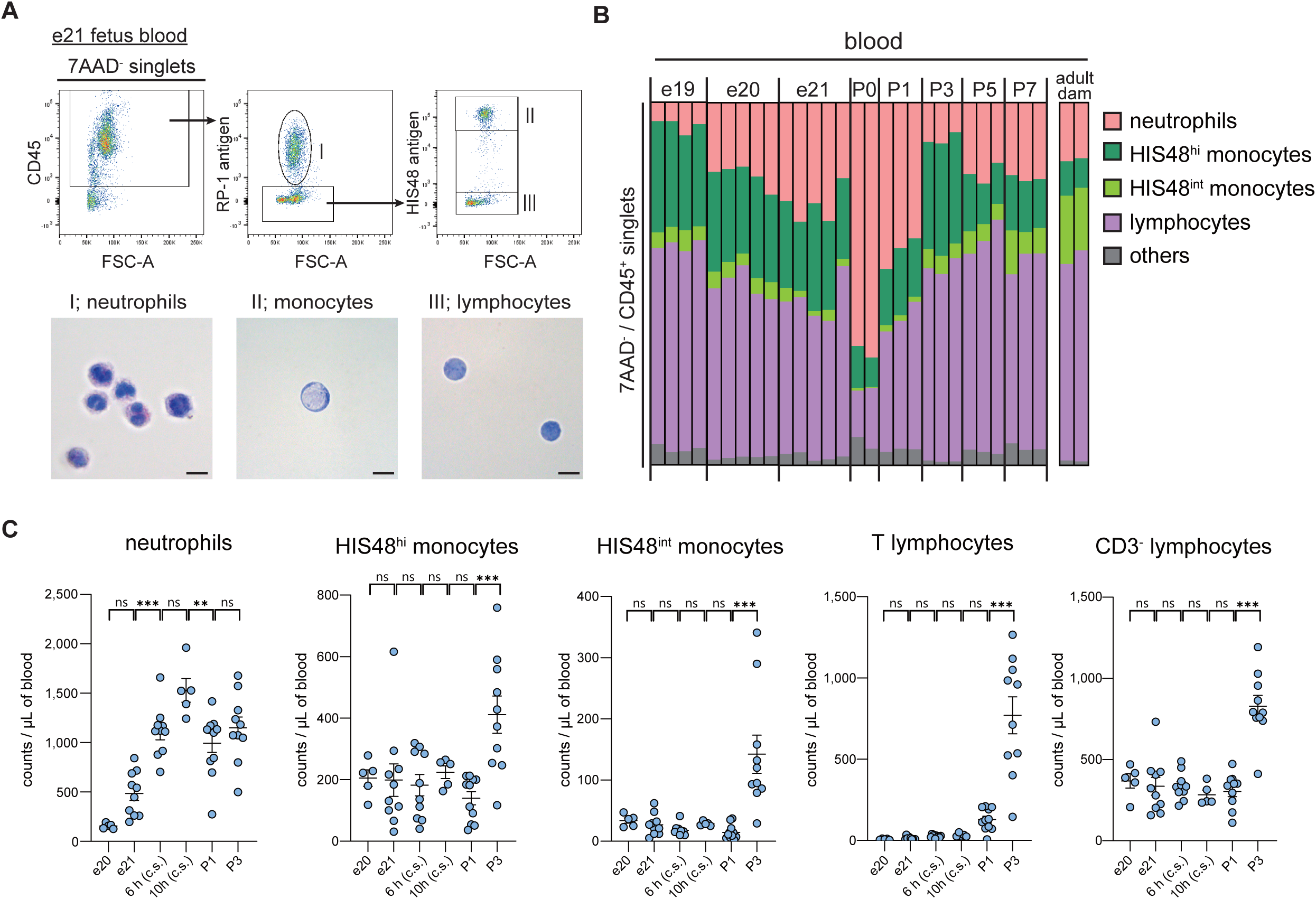
Postnatal neutrophil surge was observed in rat neonates. (A) Flow cytometry and cell sorting analysis of blood leukocytes from an e21 rat fetus. CD45^+^ / RP-1^+^ cells (neutrophils), CD45^+^ / RP-1^−^ / HIS48^hi^ cells (monocytes), and CD45^+^ / RP-1^−^ / HIS48^−^ cells (lymphocytes) were sorted and stained with Wright-Giemsa stain. Scale bars; 10 μm. (B) The proportions of blood leukocytes (CD45^+^ cells) of rat fetuses and neonates at e19, e20, e21, P0 (6-12 h after birth), P1 (12-24 h after birth), P3, P5, and P7. The proportions of blood leukocytes of adult rats (dams) are also shown. Each column represents a single fetus, neonate, or dam, except for the e19 and e20 samples. For e19 and e20, blood samples from two or three fetuses were pooled as one sample. (C) Changes in absolute counts per blood volume for neutrophils, HIS48^hi^ monocytes, HIS48^int^ monocytes, T lymphocytes, and CD3^−^ lymphocytes from e20 to P3. Neonates at 6 h or 10 h were delivered by caesarian section (c.s.). *n* = 5-11 from one or two independent dams for each group. ***, *P* < 0.001 by one-way ANOVA followed by Tukey’s multiple comparison tests. ns, not significant.

Subsequently, we investigated the postnatal neutrophil surge in rat neonates. The proportion of blood neutrophils exhibited an abrupt increase within 6-12 hours after birth (P0), followed by a decline toward P1 (12-24 hours) and P3 (48-72 hours) (Fig. 1B). Subsequently, the absolute count of each cell type per blood volume was determined using fluorescent counting beads (Fig. S3A). Additionally, an anti-CD3 antibody was employed to ascertain the number of T lymphocytes (Fig. S3B and C). Figure 1C illustrates the enumeration of each cellular population at e20 or e21, 6 h or 10 h postpartum (cesarean delivery), P1, and P3 (vaginal delivery). The number of neutrophils in the blood increased from 484/μL at e21, to 1,109/μL at 6 h, then to 1,527/μL at 10 h on average. In contrast, the number of other cell populations remained constant throughout this period. The count of monocytes and lymphocytes increased abruptly from P1 to P3 and steadily from P3 to P7 (Fig. S4). These data demonstrate, for the first time, that the postnatal neutrophil surge also occurs in rat neonates.

### The postnatal neutrophil surge was not attributable to the bone marrow neutrophil pool

The number of circulating neutrophils is maintained by a balance of neutrophil proliferation, storage, and release (12). In adults, a surge in circulating neutrophils is caused by emergency granulopoiesis in the bone marrow. In the event of a systemic bacterial infection, emergency granulopoiesis occurs, which is characterized by an enhanced proliferation of myeloid progenitors and the release of immature neutrophils into the circulation (9). We thus sought to determine whether the bone marrow emergency granulopoiesis occurs in response to birth. To analyze the bone marrow of rats, we labeled erythroblasts with an anti-CD71 antibody (13) to exclude them from the analysis (Fig. S5A and B). Fig. 2A and B illustrate the representative flow cytometry plots and the composition of bone marrow cells at e21, 6 h, P1, and P3. The proportion of neutrophils in the bone marrow remained consistent from e21 to 6 h and subsequently increased at P1. Given that the postnatal neutrophil surge was already evident at 6 h of birth (Fig. 1C), the surge was not attributable to neutrophil egress from the bone marrow.

**Figure 2.**
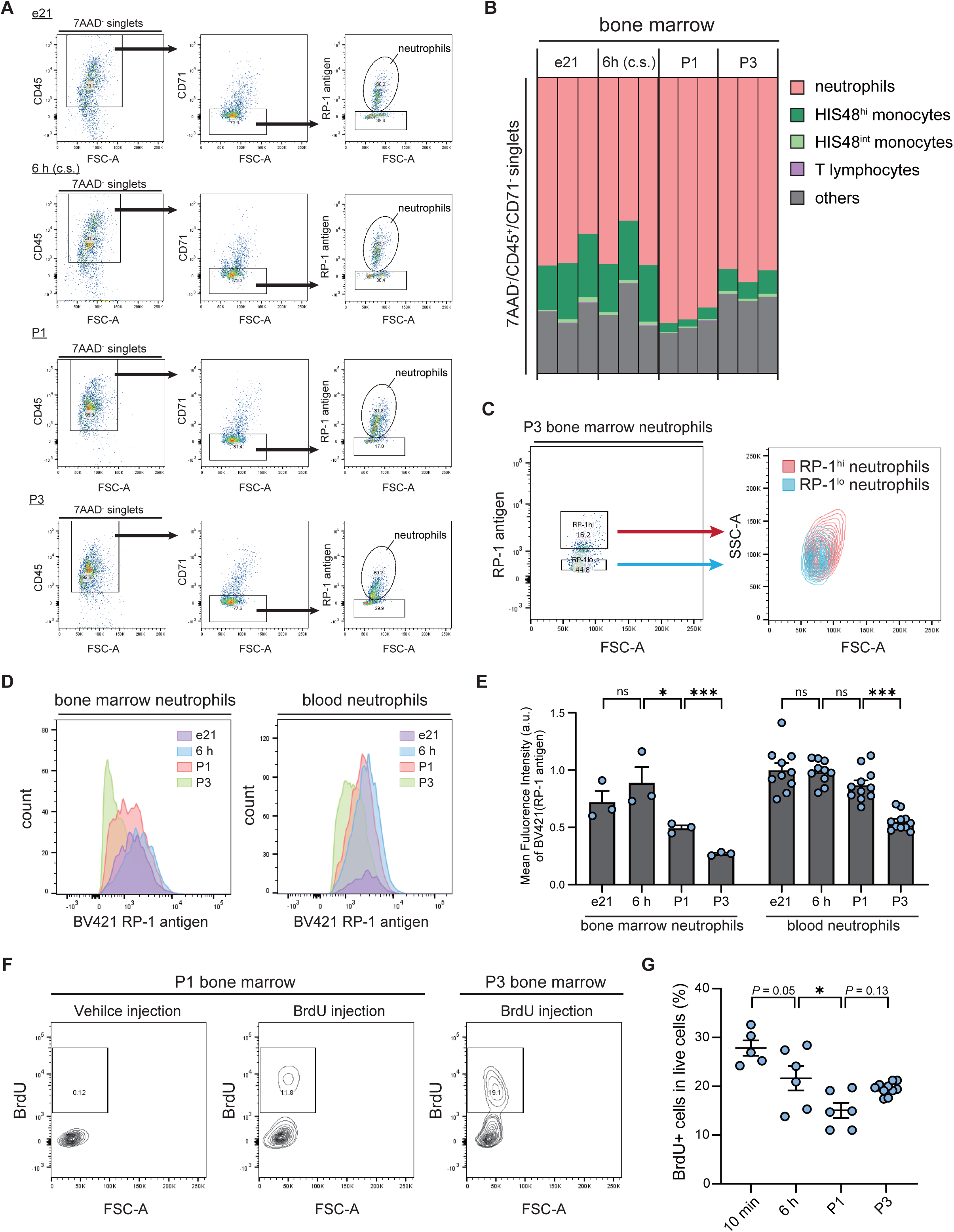
Analysis of bone marrow neutrophils in rat fetuses and neonates. (A) Representative flow cytometry plots and (B) the proportions of rat bone marrow cells (CD45^+^/CD71^−^ cells) from e21 fetus, 6 h, P1, and P3 neonates. Neonates at 6 h were delivered by caesarian section (c.s.). Each column represents a pooled sample from two individual fetuses or neonates. (C) FSC-SSC plots and of RP-1^hi^ neutrophils and RP-1^lo^ neutrophils in the bone marrow of the P3 rat neonates. (D) Representative histograms and (E) the mean fluorescence intensity of BV421 (RP-1 antigen) of the bone marrow neutrophils and the blood neutrophils from e21, 6 h, P1, and P3. The mean fluorescence intensity was normalized by the mean value of the e21 bone marrow neutrophils. Bone marrow neutrophils; *n* = 3 from two individual experiments. Blood neutrophils; *n* = 10-11 from four individual experiments. *, *P* < 0.05; ***, *P* < 0.001 by one-way ANOVA followed by Tukey’s multiple comparison tests. ns, not significant. (F) Representative flow cytometry plots and (G) the proportion of BrdU^+^ cells in the bone marrows from 10 min, 6 h, P1, and P3 neonates. Neonates at 10 min and 6 h were delivered by caesarian section (c.s.). Each point represents a pooled sample from two individual neonates. *N* = 5-11 from two independent dams for each group. *, *P* < 0.05 by one-way ANOVA followed by Tukey’s multiple comparison tests.

It was observed that the bone marrow neutrophils exhibited a skewing towards low RP-1 signal intensity for P1 and P3 (Fig. 2A). The RP-1^lo^ neutrophils exhibited lower SSC profile than the RP-1^hi^ counterparts (Fig. 2C). Given that the intensity of SSC is indicative of neutrophil granularity (14), it may be inferred that the RP-1 signal intensity is reflective of neutrophil maturity. The mean RP-1 signal intensity in the bone marrow neutrophils remained consistent from e21 to 6 h but decreased toward P1 and P3 (Fig. 2D and E). A similar pattern was observed in the blood neutrophils, although the RP-1 signal intensity did not significantly decline on P1. This suggests that the bone marrow granulopoiesis was activated from P1 to P3, resulting in the release of immature neutrophils into the circulation by P3.

The observed increase in immature neutrophils in the blood indicated that emergency granulopoiesis occurred after P1. To analyze the proliferative state of bone marrow cells, we employed a bromodeoxyuridine (BrdU) injection to label proliferative cells, collected the bone marrow cells after 90 minutes, and performed flow cytometry (Fig. 2F and G). A transient decrease in the proportion of BrdU^+^ cells was observed from birth to P1, followed by a slight increase from P1 to P3. As the RP-1^+^ neutrophils were negative for BrdU (data not shown), the transient decrease on P1 may be attributed to the observed increase in the neutrophil proportion (Fig. 2B). Given the significant increase in blood lymphocyte and monocyte counts observed from P1 to P3 (Fig. 1C), and the recent evidence indicating that the bone marrow supports the expansion of hematopoietic stem cells only after birth (15, 16), the observed increase in BrdU+ cell proportion from P1 to P3 is likely indicative of a rapid expansion of hematopoietic stem cells and activation of hematopoiesis. This is distinct from the adult emergency granulopoiesis in which neutrophil production is promoted at the expense of lymphocyte production (9).

In conclusion, the dynamic changes in the proportion and maturity of neutrophils observed in the bone marrow occurred only after P1 at which point the postnatal neutrophil surge has already peaked out. Therefore, it is not reasonable to attribute the postnatal neutrophil surge, which had peaked out before P1, to the egress or activated production of bone marrow neutrophils.

### A prenatal accumulation and postnatal reduction of the neutrophil pool were observed in the rat liver

We then suspected that the postnatal neutrophil surge arises from extramedullary neutrophil pools. The fetal hematopoiesis system is well conceptualized from the mouse studies; during the fetal development, the site for hematopoiesis shifts from the yolk sac to the aorta-gonad-mesonephros region, the lymph nodes, and then to the spleen and the liver (17). In adult mice, the lungs also serve as neutrophil reservoirs, releasing these cells at the onset of infection (18, 19).

Immunohistochemistry for myeloperoxidase (MPO), a neutrophil marker, was performed on putative neutrophil reservoirs, namely the lung, spleen, and liver (Fig. 3A). The lungs of adult rats exhibited a high density of MPO-positive cells, whereas fetal and neonatal lungs displayed only a sparse presence of these cells. The density of neutrophils was high in the spleen and liver of an e21 fetus, exhibiting a notable decline following birth (Fig. 3B). The density of the MPO-positive cells in the left lateral lobe of the liver was higher than that in the right lateral lobe and decreased already at 6 h after caesarean delivery. Additionally, a perivascular accumulation of MPO-positive cells was observed in the neonatal liver (Fig. 3A), indicative of neutrophil egress after birth. It is noteworthy that the wet weight of the liver decreased after birth, while the spleen weight steadily increased from e21 to P3 (Fig. 3C).

**Figure 3.**
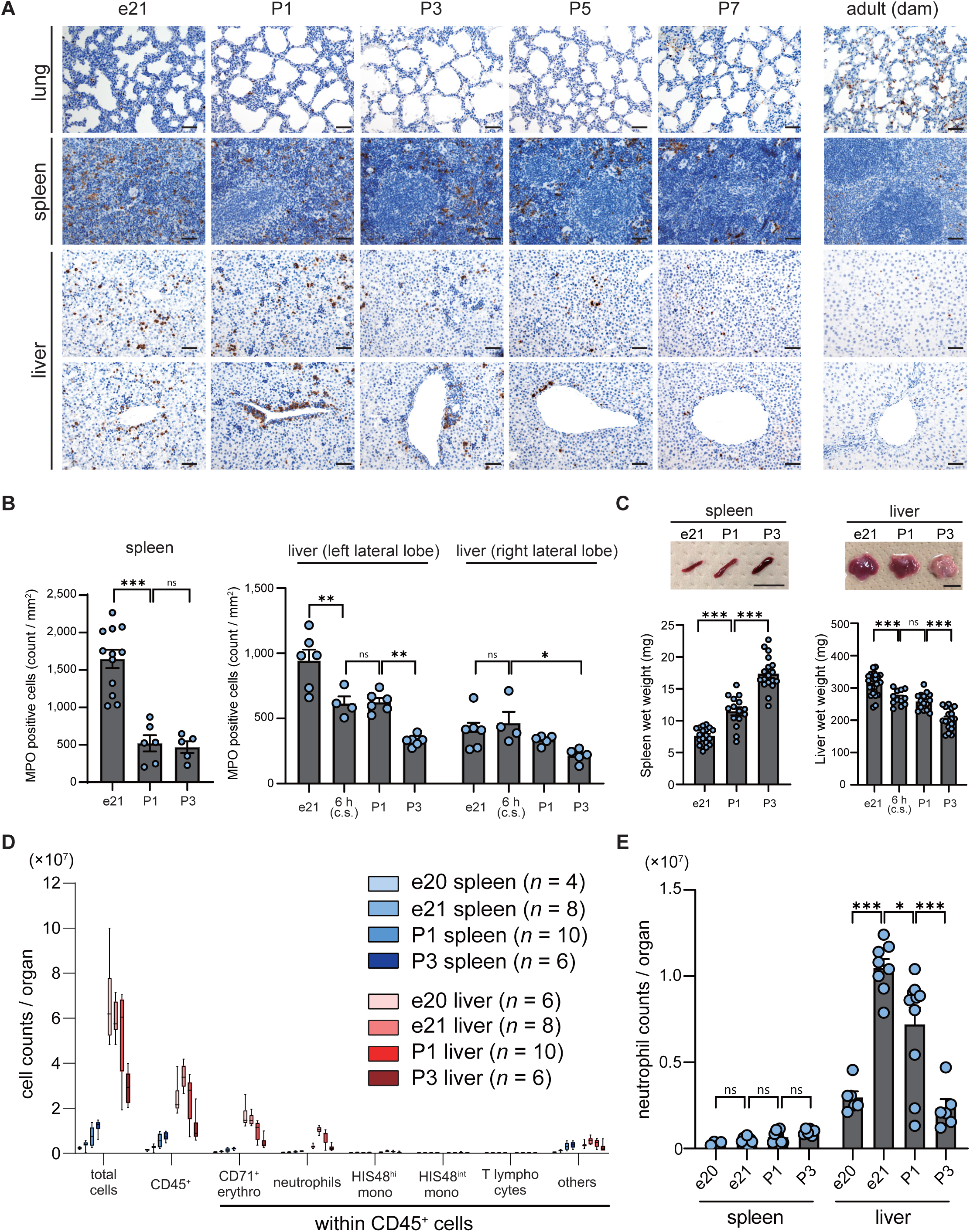
Postnatal reduction of neutrophils was observed in the liver of rat neonates. (A) Representative images of immunohistochemistry for myeloperoxidase (MPO) in rat organs (lung, spleen, and liver) at e21, P1, P3, P5, P7, and rat dam. Scale bars; 50 μm. (B) The MPO-positive cell counts/area of rat spleen at e21, P1, or P3, and of liver (left lobe or right lobe) at e21, 6h, P1, or P3. spleen; *n* = 5-12, liver; *n* = 4-6 from two independent dams for each group. (C) The representative images and the wet weights of rat spleen and liver at e21, P1, or P3. Scale bars; 10 mm. spleen; *n* = 15-18, liver; *n* = 12-28 from two or three independent dams for each group. (D, E) Whole organ flow cytometry analysis of rat spleen and liver at e20, e21, P1, and P3. (D) Total cell counts were determined by the cell counting of a cell suspension obtained from a whole spleen or liver. Cell counts of each cell population per single spleen or liver were estimated from the total cell counts and the flow cytometry data. *n* = 4-10 from one or two independent dams for each group. (E) The data of neutrophil counts in (D) were detailed. *, *P* < 0.05; **, *P* < 0.01; ***, *P* < 0.001 by one-way ANOVA followed by Tukey’s multiple comparison tests. ns, not significant.

Subsequently, a cell suspension was prepared from a whole spleen or a whole liver, and flow cytometry was performed to estimate the neutrophil count per organ (Fig. S6). Fig. 3D provides a summary of the cell count for each cell fraction in the spleens and livers from e20 to P3, with Fig. 3E highlighting the neutrophil count. As observed in the organ weight data (Fig. 3C), the total cell counts in the spleen exhibited a continuous increase from e20 to P3, accompanied by a slight increase in the neutrophil count (Fig. 3E). In contrast, the total cell counts of the liver exhibited a decline following birth, largely attributable to a reduction in the number of CD71^+^ cells (erythroblasts) and neutrophils (Fig. 3D). The livers contained a significantly greater number of neutrophils than the spleens. The liver neutrophil count decreased from 1.05×10L on e21 to 0.72×10L on P1, then to 2.36×10L on P3 (Fig. 3E). During the final day of gestation (from e20 to e21), a 3.53-fold increase in the liver neutrophil count was observed, while the erythroblast count remained unchanged. These findings suggest that neutrophils accumulate in the fetal liver during late gestation and are released into the circulation after birth.

### Analysis of Lys-EGFP mouse neonates supported the liver-origin of the postnatal neutrophil *surge*

Subsequently, we aimed to observe the postnatal reduction of liver neutrophils in individual neonates. To this end, we observed *Lys-EGFP* mice, in which the EGFP (enhanced green fluorescent protein) gene has been inserted into the lysozyme M (*Lys*) gene locus. In these mice, EGFP is highly expressed in neutrophils and to a lesser extent in monocytes (20). The *in vivo* imaging system (IVIS) was employed to observe caesarean-delivered full-term (e18) neonates (Fig. 4A and B). EGFP signals were detected in the upper abdomen region (liver ROI) of the neonates, which decreased over time through 720-750 min after birth, thereby supporting the liver origin of the postnatal neutrophil surge. It is noteworthy that the signal intensity in the oral region (oral ROI) exhibited an increase over time (Fig. 4A and B), suggesting that the neutrophils may have migrated into oral tissues. Additionally, the P1, P2, and P3 neonates were observed (Fig. 4C). The pups were delivered vaginally and nursed by the dam. Autofluorescence signals were observed in the stomach and intestine of breastfed wild-type neonates, but not in other organs, including the liver (Fig. S7A). Given the detection of an autofluorescence signal in the urine of both *Lys-EGFP* and wild-type neonates (Fig. S7B and C), the urine was removed by bladder puncture prior to observation. Upon examination of the dissected organs, a 30-minute *Lys-EGFP* neonate exhibited robust EGFP signals in the liver, spleen, and femur bone marrow (Fig. 4C). EGFP signals in the sternum bone marrow were only detected after P1 and were more evident on P2 and P3. These results corroborate those of the rat experiments, indicating a postnatal reduction of liver neutrophils before P1 and the activation of bone marrow granulopoiesis from P1 to P3.

**Figure 4.**
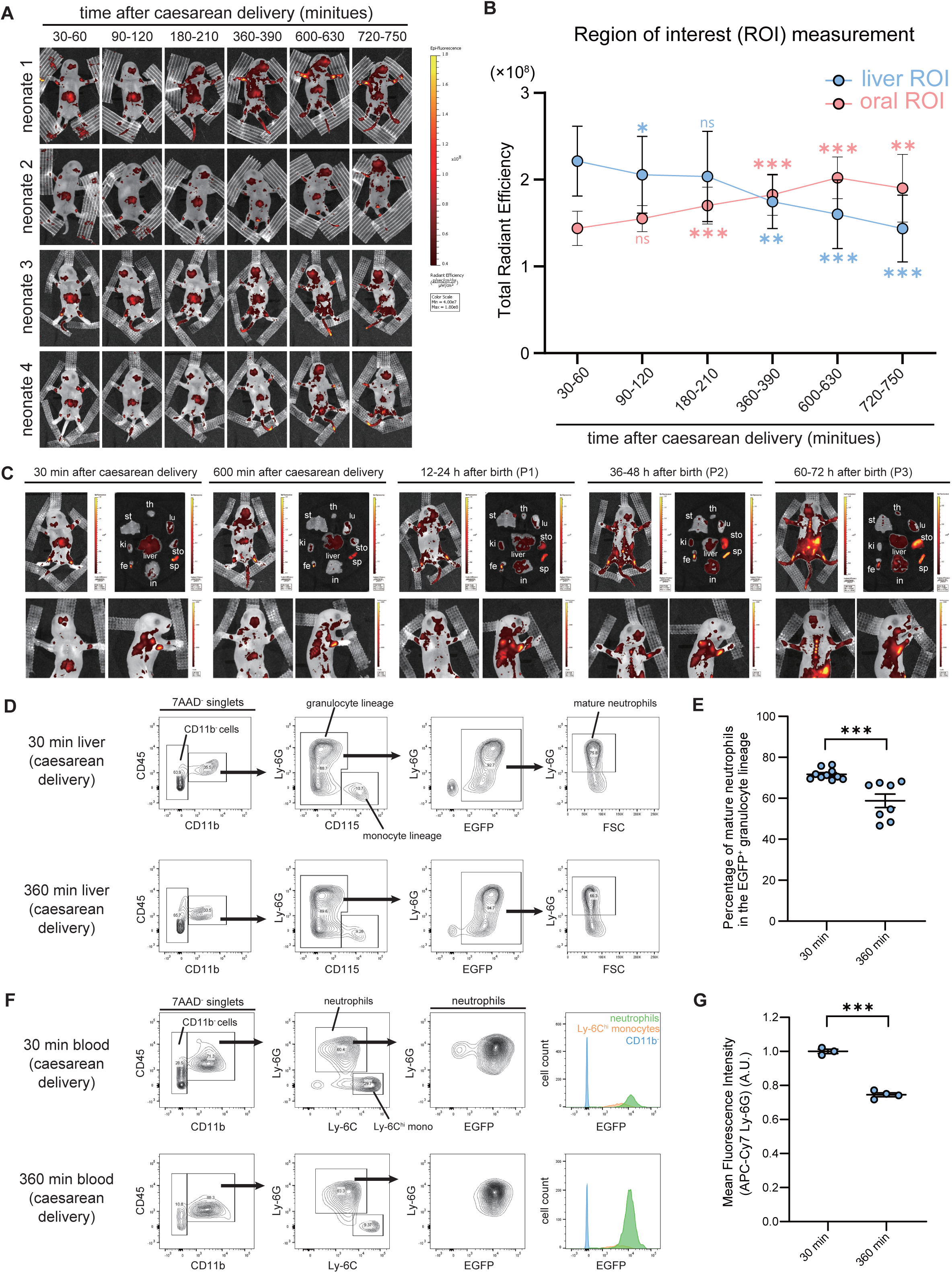
*Lys-EGFP* mouse neonates exhibited postnatal reduction of liver neutrophil pool. (A) Representative fluorescence images obtained by IVIS. *Lys-EGFP* mouse fetuses were delivered at e18 by caesarean section. Each row represents an individual neonate imaged over time up to 720-750 min after birth. (B) Florescence intensity of (A) was measured at the liver region of interest (ROI) and the oral ROI. mean ± S.D. *n* = 10 from two individual dams. *, *P* < 0.05; **, *P* < 0.01 ***, *P* < 0.001 vs. 30-60 min group of each ROI by repeated measures one-way ANOVA followed by Dunnett’s multiple comparisons test. ns, not significant. (C) Representative fluorescence images of *Lys-EGFP* mouse neonates at 30 min, 600 min, P1, P2, and P3 of birth. Whole body image (upper left), dissected organs (upper right), zoomed images of the upper body (lower panels) are shown for each group. th: thymus, st: sternum, lu: lung, ki: kidney, sto: stomach, sp: spleen, fe: femur, in: intestine. (D and F) Representative flow cytometry plots of *Lys-EGFP* neonatal liver (D) and blood (F) at 30 min and 360 min after caesarean delivery. (E) Comparison of the proportion of mature neutrophils in (D). *n* = 8-10 from two independent dams. (G) Comparison of mean fluorescence intensity of APC-Cy Ly-6G in the blood neutrophils in (F). Each point represents a pooled sample from two or three individual neonates. *n* = 3-4 from two independent dams. ***, *P* < 0.001 by unpaired Student’s *t*-test.

Subsequently, a flow cytometric analysis was conducted on the *Lys-EGFP* neonatal liver (Fig. 4D). Myeloid cells were identified as CD45^+^/CD11b^+^; then monocyte-lineage cells were designated as Ly-6G^−^/CD115^+^ (21). The majority of CD115^−^ cells exhibited EGFP positivity (Fig. 4D), thereby indicating that this fraction represents the granulocyte lineage. Additionally, monocyte lineage cells exhibited EGFP expression, albeit with significantly lower signal intensity and cell count compared to the granulocyte lineage (Fig. S8A and B). This suggests that the EGFP signal in the liver is predominantly derived from the granulocyte lineage. Within the granulocyte lineage, we detected Ly-6G^hi^ mature neutrophils (22) (Fig. 4D). Cell sorting and Wright-Giemsa staining were employed to confirm that the CD45^+^/CD11b^+^/CD115^+^ cells are morphologically monocytes and that the CD45^+^/CD11b^+^/CD115^−^/EGFP^+^/Ly-6G^hi^ cells are mature neutrophils with a segmented nucleus (Fig. S8C). The CD45^+^/CD11b^+^/CD115^−^/EGFP^+^/Ly-6G^lo^ fraction included cells with a spherical nucleus or with a ring-shaped nucleus, which are characteristics of immature neutrophils (23). No evidence of eosinophilic or basophilic cells was found in this fraction. In the livers, the proportion of mature neutrophils within the granulocyte population exhibited a significant decline from 30 to 360 minutes postpartum (Fig. 4E), suggesting that the mature neutrophils were undergoing selective release. Additionally, the observation of cryosections of the *Lys-EGFP* neonatal livers also demonstrated a perivascular accumulation of EGFP^+^ cells after birth, indicative of neutrophil emigration (Fig. S9).

The analysis of Lys-EGFP neonatal blood revealed a notable elevation in the proportion of neutrophils (CD45^+^/CD11b^+^/Ly-6G^+^) at 360 minutes postnatal (Fig. 4F), indicative of the postnatal neutrophil surge. The Ly-6G^+^ blood neutrophil population exhibited characteristic changes from 30 min to 360 min postpartum. At 30 min, the blood contained EGFP^lo^ neutrophils, which were undetectable at 360 min. Additionally, there was a decrease in membrane Ly-6G expression from 30 min to 360 min. As the Ly-6G^+^ neutrophils in the livers were predominantly EGFP^hi^ and that the liver contained neutrophils with various levels of Ly-6G membrane expression (Fig. 4D), it can be postulated that these observed changes may be due to the predominance of the liver-derived neutrophils in the blood. It is noteworthy that the neonatal blood sample taken at 360 minutes after birth did not contain a detectable level of Ly-6C^lo^ (non-classical) monocytes, which was distinctive from adult blood (Fig. 4F and Fig. S10).

### Neutrophil-biased myeloid differentiation was promoted in a G-CSF-dependent manner in the fetal mouse liver during late gestation

The presence of Ly-6G^lo^ immature granulocytes in the mouse neonatal liver indicated that granulopoiesis is active in the fetal liver at the time of birth. We also observed the significant increase in the number of neutrophils in the rat fetal liver from e20 to e21 (Fig. 3E). From these data, we hypothesized that the granulopoiesis is activated in the fetal liver during the late gestation, resulting in the accumulation of the liver neutrophil pool toward birth.

Neutrophils are derived from granulocyte-monocyte progenitors (GMP), which also give rise to monocytes; GMP are derived from common myeloid progenitors (CMP), which also give rise to megakaryocyte-erythrocyte progenitors (MEP) (24). We first examined the proliferative state of these progenitors by flow cytometry. We injected BrdU into wild-type pregnant mice and analyzed the fetal liver (Fig. 5A). BrdU was successfully incorporated into fetal liver cells and was specifically detected by the anti-BrdU antibody (Fig. 5B). From e14 to e16 and to e18, the proportion of GMP and CMP decreased significantly (Fig. 5C and D), accompanied by a decrease in the BrdU-positive rate (Fig. 5E). Instead, we observed a significant increase in differentiated leukocytes (CD45^+^/Lin^+^). These data suggest that proliferation of myeloid precursors in the fetal liver was arrested around birth, while leukocyte differentiation continued.

**Figure 5.**
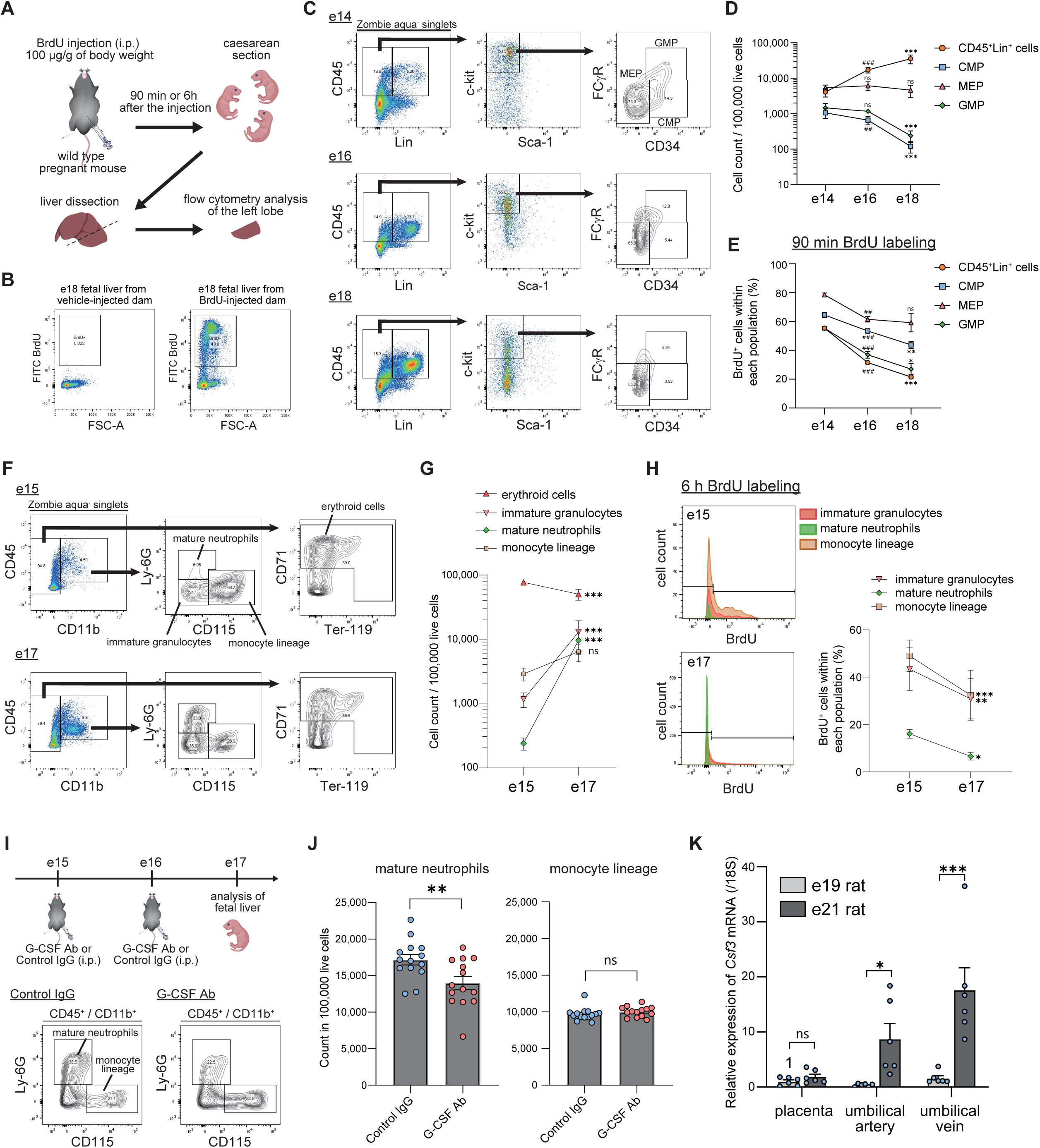
Neutrophil-biased myeloid differentiation was observed in the mouse fetal liver during the late gestational period. (A) Experimental scheme of BrdU treatment and flow cytometry for fetal liver. (B) Representative flow cytometry plots for fetal livers collected from vehicle-injected dam or BrdU-injected dam. (C) Representative flow cytometry plots for CMP, MEP, and GMP in the fetal livers collected from BrdU-injected pregnant mice at e14, e16, or e18. (D) Changes in the cell counts and (E) the BrdU positive rate in (C). (B-E) Caesarean section was performed after 90 min of BrdU injection. *n* = 8-10 from two independent dams for each group. ##, *P* < 0.01; ###, *P* < 0.001 (vs. e14) and *, *P* < 0.05; **, *P* < 0.01; ***, *P* < 0.001 (vs. e16) by one-way ANOVA followed by Tukey’s multiple comparison tests. ns, not significant. (F) Representative flow cytometry plots of fetal livers collected from pregnant mice at e15 or e17. (G) Changes in the cell counts and (H) representative histograms and the BrdU positive rates in (F). (F-H) Caesarean section was performed after 90 min of BrdU injection. *n* = 8-9 from two independent dams for each group. ***, *P* < 0.001 vs. e15 by two-way ANOVA followed by Sidak’s multiple comparisons test. (I) Experimental scheme of G-CSF neutralizing antibody treatment and the representative flow cytometry plots of the fetal livers at e17. (J) Cell counts of neutrophils and monocyte lineage in (J). *n* = 14 from three independent dams for each group. **, *P* < 0.01 by unpaired Student’s *t*-test. ns, not significant. (K) Relative mRNA expression of *Csf3* (G-CSF) gene in the placenta, umbilical artery, and umbilical vein from rat fetuses on e19 or e21. *n* = 6 from two individual dams for each group. *, *P* < 0.05; ***, *P* < 0.001 by two-way ANOVA followed by Sidak’s multiple comparisons test.

We then examined the myeloid population in the fetal liver at e15 and e17 (Fig. 5F and G). Erythroid cells were detected as CD71^+^ and/or Ter-119^+^ within the CD11b^−^ cells (21). The proportion of immature granulocytes (CD45^+^/CD11b^+^/CD115^−^/Ly-6G^lo^) and mature neutrophils (CD45^+^/CD11b^+^/CD115^−^/Ly-6G^hi^) increased significantly from e15 to e17, reversing that of the monocyte lineage (CD45^+^/CD11b^+^/CD115^+^/Ly-6G^−^) (Fig. 5G). At e15, immature granulocytes were approximately 45% positive for BrdU after 6 h of BrdU labeling, whereas mature neutrophils were only 18% positive (Fig. 5H), supporting that the Ly-6G^lo^ fraction represents immature neutrophils (23). The BrdU positive rate decreased from e15 to e17 in both immature granulocytes and in mature neutrophils (Fig. 5H), suggesting that the neutrophil differentiation or maturation rather than the granulocyte proliferation was promoted during late gestation.

Next, we sought to determine factors mediating the neutrophil-biased myeloid differentiation in the fetal liver. Granulocyte-colony stimulating factor (G-CSF) is a potent inducer of neutrophil differentiation (25). To investigate the contribution of G-CSF, we injected wild-type pregnant mice with G-CSF-neutralizing antibodies on e15 and e16 and analyzed the fetal livers on e17 (Fig. 5I). Compared with control IgG administration, G-CSF neutralizing antibody significantly decreased the proportion of mature neutrophils, while the monocyte lineage was not affected (Fig. 5J), indicating that the neutrophil-biased differentiation was partially mediated by G-CSF.

We also investigated possible sources of G-CSF in the late gestational period. In full-term fetal rat livers (e21), the mRNA expression level of the *Csf3* (G-CSF) gene was below the detection limit by qRT-PCR (data not shown). Therefore, we hypothesized the existence of an endocrine mechanism. The placenta serves as an endocrine organ and shows a distinctive secretion repertoire depending on gestational age (26); G-CSF is reported to be expressed in human placenta (27). In addition, the *CSF3* gene is highly expressed in endothelial cells according to the Human Protein ATLAS (proteinatlas.org). Therefore, we examined the mRNA expression of *Csf3* in rat placenta and umbilical blood vessels at late gestation (e19 and e21) (Fig. 5K). From e19 to e21, the *Csf3* expression did not change in the placentas, but significantly increased in the umbilical arteries and in the umbilical veins. These data suggest that umbilical-derived G-CSF may promote neutrophil-biased myeloid differentiation in the fetal liver.

### Mouse liver neutrophils showed changes in transcriptomic profiles within 3 h after birth

We then sought to determine whether liver neutrophils undergo characteristic changes in response to birth that may be relevant to the neutrophil egress after birth. To this end, we performed bulk RNA sequence analysis (RNA-seq) on neutrophils collected from *Lys-EGFP* mouse neonates (Fig. 6A). Liver neutrophils were collected by FACS immediately after caesarean delivery (pre), at 10 min (confirmed spontaneous breathing), or at 3 h after the caesarean delivery. A total of 6 samples from 4 dams were subjected to RNA-seq. Fig. 6B is the result of Principle Component Analysis (PCA), showing that liver neutrophils exhibit transcriptomic changes from 10 min to 3 h, which were commonly observed between the two independent dams. We then compared the 10 min group and the 3 h group in detail. When we set the cutoff *P* value at 10^−12^, we detected 47 genes that were differentially expressed between 10 min and 3 h, of which 43 genes were upregulated after birth (Fig. 6C and D). The most higly expressed genes included genes such as *S100a8*, *S100a9*, *Lcn2*, and *Ly6G* (Fig. 6E), which are transcriptional signatures that define neutrophils (28). Gene set enrichment analysis (GSEA) showed that genes related to neutrophil degranulation were either up- or down-regulated after birth (Fig. 6F). Genes related to neutrophil activation (signaling by Rho GTPase and signaling by VEGF) (29, 30) and to platelet activation tended to be downregulated after birth (Fig. 6F). Genes related to focal adhesion were also downregulated, which may be related to the release of liver neutrophils. Taken together, liver neutrophils showed transcriptional changes after birth.

**Figure 6.**
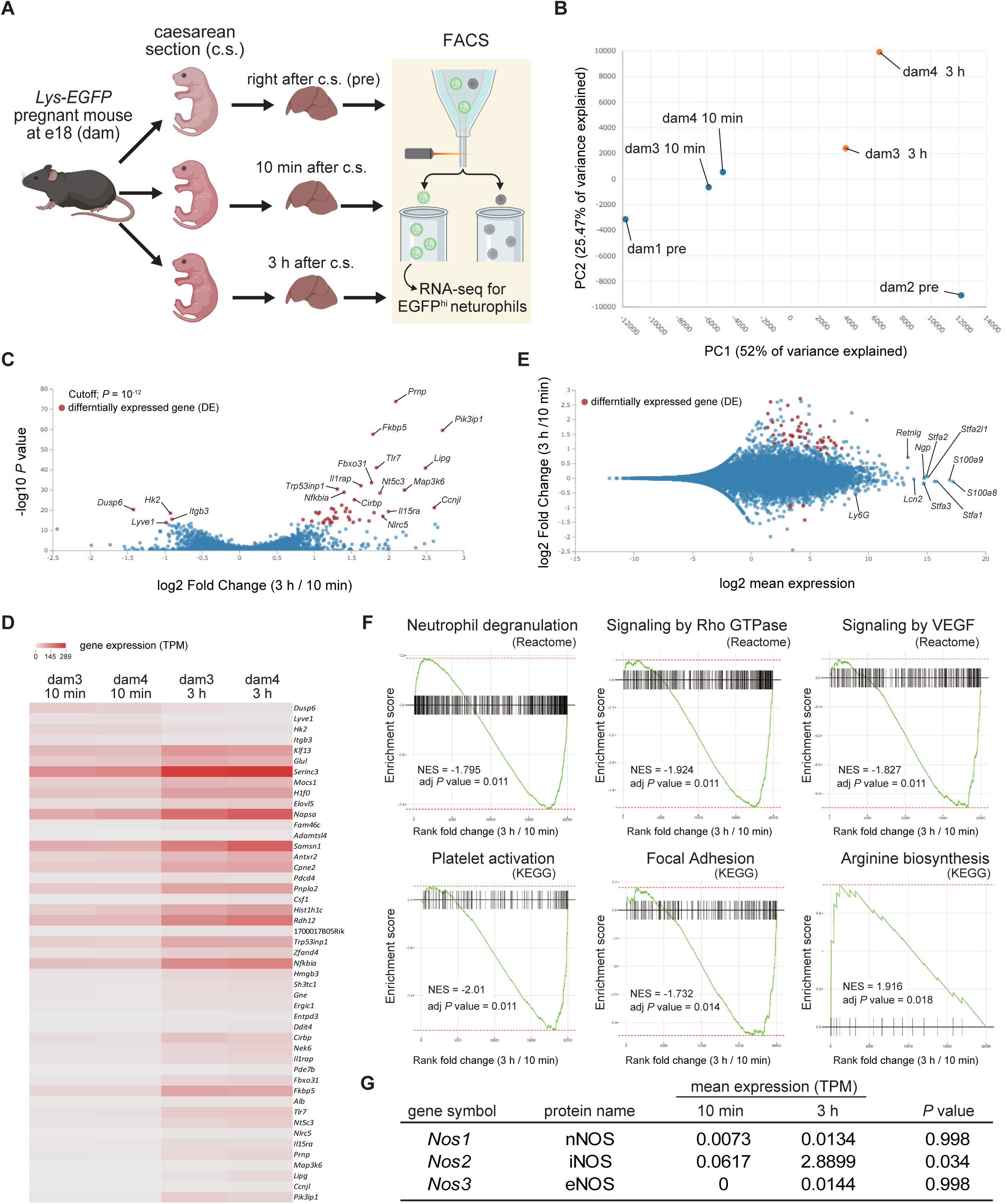
Liver neutrophils exhibited changes in transcriptome profiles within 3 h after birth. (A) Experimental scheme showing neutrophil collection from *Lys-EGFP* neonates. (B) Principal component analysis (PCA) plot for RNA-seq data of liver neutrophils obtained right after the caesarean section (pre), 10 min after the c.s., or 3 h after the c.s. Data were obtained from four dams (dam1-4). (C) Volcano plot showing the differentially expressed genes (DE) between the 10 min group and the 3 h group. Cutoff *P* value is set at 10^−12^. (D) Heatmap showing the gene expressions of DE in TPM (transcripts per million). (E) MA plot of the RNA-seq data. Red dots correspond to the DE in (C). (F) Gene set enrichment analysis (GSEA) plots of the RNA-seq data. Each plot shows the pathway name, normalized enrichment score (NES), and the adjusted *P* value. (G) Comparison of the expressions of *Nos* subtypes between the 10 min and the 3 h liver neutrophils.

### NOS inhibitors inhibited the postnatal neutrophil surge in rat neonates

Finally, we sought to determine the stimuli responsible for the release of liver neutrophils following birth. Nitric oxide (NO) is produced by the catalysis of arginine, which is mediated by nitric oxide synthase (NOS). NO modulates the functions of neutrophils in a variety of ways, including the promotion of neutrophil migration (31, 32). Kawamura *et al.* demonstrated that the mRNA expression of inducible NOS (iNOS, *Nos2*) was transiently upregulated in the mouse neonatal liver leukocytes at 3-6 hours after birth (33). Our RNA-seq results also demonstrated that genes associated with arginine biosynthesis and *Nos2* were upregulated in liver neutrophils following birth, while *Nos1* (nNOS) and *Nos3* (eNOS) were barely detected (Fig. 6F and G). We therefore postulated that the postnatal upregulation of iNOS is responsible for the egress of neutrophils from the liver. In comparison to the other NOS isoforms, iNOS displays a considerably shorter cellular turnover period, with a half-life of 1.6 ± 0.3 hours (34). The cleavage of iNOS by calpain I results in the production of an inactive form of the protein, which can be detected by western blotting at approximately 70 kDa (35). The abundance of iNOS protein was examined in the left lateral lobes of the rat neonatal livers (Fig. 7A). The cleaved iNOS was the predominant form observed, indicating a rapid turnover. Following the birth of the rats, a slight increase in the full-length iNOS was observed, accompanied by a significant and transient increase in the cleaved form (Fig. 7B). Subsequently, we investigated whether the administration of a NOS inhibitor could mitigate the postnatal neutrophil surge in rat neonates (Fig. 7C). The administration of a non-selective NOS inhibitor (L-NAME) and an iNOS-selective inhibitor (1400W) both resulted in the inhibition of the extent of the postnatal neutrophil surge at 6 h after caesarean delivery, accompanied by the retention of liver neutrophils (Fig. 7D). These results indicate that the upregulation of iNOS is responsible for the release of liver neutrophils and consequently for the postnatal neutrophil surge.

**Figure 7.**
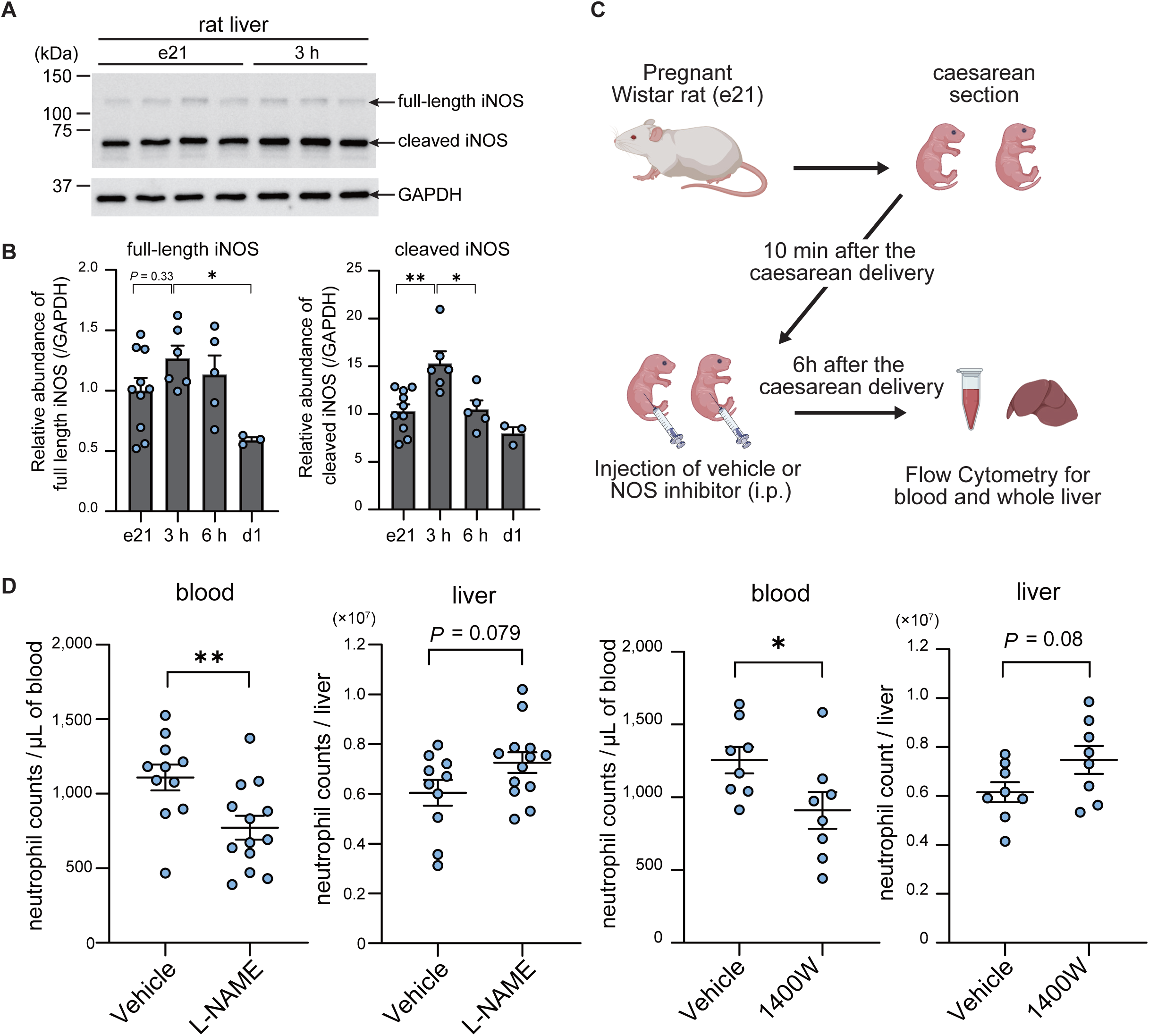
Postnatal neutrophil surge was mediated by iNOS in rat neonates. (A) Representative western blots showing iNOS and GAPDH abundance in rat neonatal liver right after caesarean section or after 3 h of caesarean delivery. (B) Relative abundance of full-length iNOS and cleaved iNOS normalized by the abundance of GAPDH in the left lateral lobes of neonatal livers at e21, 3 h, 6 h, and P1. *n* = 3-10 from two independent dams for each group. *, *P* < 0.05; **, *P* < 0.01 by one-way ANOVA followed by Tukey’s multiple comparison tests. (C) Experimental scheme showing NOS inhibitor administration for rat neonates. (D) Neutrophil counts in blood and livers from rat neonates administered with either vehicle or NOS inhibitor (L-NAME or 1400W). L-NAME; *n* = 10-13 from three independent dams for each group. 1400W; *n* = 8 from two independent dams for each group. *, *P* < 0.05; **, *P* < 0.01 by unpaired Student’s *t*-test.

## Discussion

In this study, we showed by examining rat and mouse fetuses and neonates that the postnatal neutrophil surge arises from the liver neutrophil pool. The activation of bone marrow granulopoiesis was observed only after 24-72 h of birth, at which the postnatal neutrophil surge had peaked out. Conversely, the liver neutrophils underwent a rapid decline following birth, occurring concurrently with the perivascular neutrophil accumulation in the livers and the surge of circulating neutrophils. During late gestation, myeloid cell differentiation in the fetal livers was biased toward neutrophils which was partially mediated by G-CSF. After birth, liver neutrophils underwent transcriptomic changes exemplified by the upregulation of *Nos2* (iNOS) gene. Inhibition of iNOS hindered the release of liver neutrophils and the onset of the postnatal neutrophil surge. Our data depict a previously unknown source of the postnatal neutrophil surge and a stimulus initiating the surge.

The results demonstrated that the rat is an appropriate model for investigating the postnatal neutrophil surge. The objective of the initial experiment was to ascertain whether neutrophils could be identified by flow cytometric analysis of fetal or neonatal rat blood. An anti-rat granulocyte antibody, RP-1, was observed to specifically label rat fetal and neonatal neutrophils (Fig. 1A and Fig. S1). Although HIS48 antigen is considered a marker for rat granulocytes, our results indicate that it is also expressed on monocytes. This finding aligns with previous reports that suggest the HIS48^hi^ cells are classical monocytes and the HIS48^int^ cells are non-classical monocytes (11, 36). By using RP-1, it was possible to analyze neonatal rat blood neutrophils without the necessity of pooling samples from multiple pups. This study demonstrated for the first time that the postnatal neutrophil surge is also exhibited by rats, with a similar time course to that observed in human neonates. Consequently, rats may be considered a suitable model for examining the mechanisms of the postnatal neutrophil surge.

The proportion of HIS48^int^ monocytes was observed to be low in rat neonatal blood and exhibited a gradual increase from P3 to P7 (Fig. 1B). Additionally, the proportion of non-classical monocytes (Ly-6C^lo^ monocytes) was markedly diminished in the neonatal mouse blood relative to that observed in adult mice (Fig. 4F and Fig. S10). This characteristic may be pertinent to the immunological properties of neonates during the initial week of life. Nevertheless, the physiological relevance is beyond the scope of the present study and requires further investigation.

We could not attribute the postnatal neutrophil surge to the bone marrow neutrophil pool. RP-1 has been proposed as an activation marker for rat neutrophils, given that the membrane expression of the RP-1 antigen is upregulated upon stimulation with phorbol myristate acetate (PMA) (37). The results of the present study also indicated that RP-1 can be used as a maturation marker for rat neutrophils. This led us to conclude that the activation of bone marrow granulopoiesis accompanying the increase in blood immature neutrophils occurs only after P1, at which the postnatal neutrophil surge has already peaked out. The activation of bone marrow granulopoiesis was distinct from that observed in adult emergency granulopoiesis, as evidenced by a significant increase in circulating monocytes and lymphocytes (Fig. 1C).

Our data demonstrated that neutrophils accumulate in the fetal livers prior to birth. In the prevailing view of fetal hematopoiesis, the liver plays a supportive role in the expansion of hematopoietic stem cells (HSCs), facilitating their migration to the bone marrow, so that the bone marrow starts to serves as the primary site for hematopoiesis prior to birth (38). However, recent studies have presented evidence that calls this notion into question. Ganuza *et al*. demonstrated that the contribution of fetal liver HSC to adult bone marrow hematopoiesis was minimal (16). Hall *et al.* demonstrated that murine bone marrow does not support the expansion of HSC and progenitor cells before birth (15). Yokomizo *et al.* similarly demonstrated that the contribution of fetal liver HSC to the postnatal generation of functional blood cells is minimal. Instead, newly identified pre-HSPC (pre-hematopoietic stem and progenitor cell) yielded progenitors and differentiated blood cells in a non-hierarchical manner that does not require differentiation into HSCs (39). These reports indicate that the fetal liver supports the production of functional blood cells until birth, after which hematopoiesis in the bone marrow is activated. Our data also supported this notion, as evidenced by a significant increase in the proportion of differentiated leukocytes, particularly neutrophils, during the final three days of the mouse gestational period (Fig. 5A-G).

The prenatal accumulation of liver neutrophils was indicated to be a consequence of granulopoiesis within the liver. In adults, approximately half of the circulating neutrophils are trapped and retained on the endothelium of tissue vasculature (40), forming the so-called “marginated neutrophil pool.” The liver, spleen, and lung are the primary organs containing this marginated pool (41). A recent report demonstrated that in the fetal (e14.5) and the neonatal (P0) mouse livers, Ly-6G^+^ neutrophils predominantly localize to the liver parenchyma rather than the intravascular space (42). Our results demonstrated that in the fetal (e17) and neonatal (30-360 min) livers, there was a gradient in membrane Ly-6G expression within the granulocyte lineage (Fig. 4D and Fig. 5F). Additionally, the Ly-6G^lo^ population exhibited morphologically immature neutrophils (Fig. S8C). These findings suggest that the prenatal accumulation of liver neutrophils is more likely to result from granulopoiesis within the liver rather than margination of circulating neutrophils. In this study, we demonstrated that the differentiation of neutrophils over monocytes was promoted in the fetal liver in a manner dependent on G-CSF (Fig. 5I and J). The umbilical blood vessels were identified as a potential source of G-CSF. In the rat umbilical vessels, there was a notable increase in *Csf3* (G-CSF) gene expression significantly increased from e19 to e21 (Fig. 5K), which occurred concurrently with a significant rise in liver neutrophils (Fig. 3E).

The observed difference in neutrophil density between the left and right lateral lobe of the rat fetal livers (Fig. 3B) may be attributed to the difference in blood supply. Prior to birth, the hepatic artery plays a minor role in the perfusion of the liver. The majority of venous perfusion to the fetal liver, ranging from 80 to 85%, originates from the umbilical vein, with the remaining portion derived from the portal vein (43). The left lobe of the fetal liver is primarily perfused with umbilical venous blood, whereas the right lobe is predominantly perfused with portal venous blood (43). Given the upregulation of the *Csf3* gene in the umbilical blood vessels and the higher expression in the umbilical vein (Fig. 5K), it is possible that the difference in G-CSF concentration influenced the accumulation of neutrophils between the right and left lobes of the fetal livers.

The liver origin of the neutrophil surge and the late gestational liver neutrophil accumulation demonstrated in this study would explain why the surge is hindered or absent in preterm infants and in very-low-birth-weight (VLBW) infants (7, 44). A rough comparison of gestational periods indicates that the final 2-3 days of the mouse or rat gestational period correspond to approximately 3-5 weeks of the human gestational period. In human preterm neonates (gestational age < 32 weeks), the peak level of blood neutrophil count within 12 hours of birth was approximately 4,000/μL, while full-term neonates (gestational age > 37 weeks) exhibited a mean of more than 14,000/μL (44). It is plausible that the accumulation of neutrophils in the fetal liver occurs during the final five weeks of human gestation.

The process of fetal liver granulopoiesis may be subject to regulation through a complex network of interactions involving both maternal and fetal factors. In a study conducted Stelzer *et al*., the maternal blood metabolome, proteome, and immunome were analyzed over a period of 100 days preceding the day of labor (45). The findings indicated that coordinated molecular shifts were observed two to four weeks prior to delivery, suggesting a transition from immune activation to the regulation of inflammatory responses. A Study of mouse pregnancy by Collins *et al.* demonstrated that fetal liver HSC are restricted from engaging in emergency granulopoiesis even when systemic inflammation was induced. Maternal IL-10 was identified as the factor responsible for this restriction (46). The report that maternal hypertension impedes the postnatal neutrophil surge in VLBW infants (7) also indicates that the maternal factors can affect the fetal granulopoiesis.

The present study identified G-CSF as a fetal factor that mediates the accumulation of neutrophils in the fetal liver. It is possible that stimulation of the fetal G-CSF pathway may promote the accumulation of neutrophils in the fetal liver and the subsequent postnatal neutrophil surge in preterm neonates. Such a procedure may be a potential preventive measure for neonatal sepsis. However, this measure cannot currently be assessed in rats and mice, as it is not possible to induce preterm birth in rat fetuses at e20 or in mouse fetuses at e17. To assess the effects of prenatal G-CSF stimulation, it may be necessary to examine larger animals with a longer gestational period. It should be noted that the postnatal prophylactic use of G-CSF for human neonates does not conclusively decrease the risk of neonatal sepsis (47).

It has been reported that caesarean delivery results in a reduction in the extent of the postnatal neutrophil surge in comparison to vaginal delivery (6). Given the narrow temporal window during which the postnatal neutrophil surge occurs and the fact that parturition in rats typically occurs within an average of 100 minutes (48), we employed a caesarean delivery approach for the majority of experiments conducted in this study. A limitation of the present study is that it does not examine the effect of caesarean delivery compared to vaginal delivery.

The risk of neonatal sepsis is elevated in neonates delivered via caesarean section (49). Thus, stimulating the postnatal neutrophil surge may offer a potential preventive measure. While the upstream stimuli for iNOS upregulation in the livers remain unclear, hypoxia may represent a pivotal factor. In a number of cell types, hypoxia-inducible factor (HIF)-1α has been shown to activate iNOS transcription (50). Furthermore, the postnatal neutrophil surge is enhanced in human neonates born at high altitudes (4,800 feet, approximately 1,463 meters above sea level in average) (6), indicating that hypoxia is a key activator of the postnatal neutrophil surge. While there is no existing literature on this topic, it is plausible that the left lobe of the liver is subjected to transient hypoxia due to the occlusion of oxygen-rich umbilical venous blood following birth.

It is also possible that glucocorticoids may be a contributing factor to the postnatal neutrophil surge. Our RNA-seq analysis (Fig. 6C) revealed that the most significantly upregulated genes in liver neutrophils included *Prnp* and *Fkbp5*, both of which are known to be upregulated by glucocorticoid stimulation (51, 52). The plasma cortisol level in human neonates is the highest on the day of birth, subsequently decreasing until day3 (53). The concentration of cortisol was observed to be lower in neonates delivered via caesarean section (54). Additionally, glucocorticoids have been demonstrated to enhance neutrophil survival (55). Our future objective is to investigate whether hypoxia and glucocorticoids are accountable for the stimulation of the postnatal neutrophil surge.

Despite the postnatal neutrophil surge being regarded as a defensive mechanism against infection, the specific roles it plays remain unclear. Our observations of *Lys-EGFP* mouse neonates indicated that neutrophils accumulate in the oral area at the time of the postnatal neutrophil surge (Fig. 4A and B). The oral tissue section was observed, and neutrophils were detected in the oral mucosa (data not shown). In a detailed examination of the microbiota of human neonates, Ferretti and colleagues discovered that the oral microbiota is highly diverse at day1 of birth. By day3, this microbiota undergoes rapid selection to form an oral microbial niche (56). Given the concurrence of the postnatal neutrophil surge and the oral accumulation of neutrophils, it is reasonable to hypothesize that the liver-derived neutrophils participate in the host-microbe interaction in the oral tissue. It is also our objective to examine this possibility.

In summary, the presenting study revealed the previously unknown source of the postnatal neutrophil surge and set the basis for investigating the mechanisms of the surge. By further studying the mechanisms and the specific roles of the postnatal neutrophil surge, we may be able to develop preventive measures for neonatal sepsis.

## Materials and methods

### Animal experiments

We performed the animal studies under the approval of the institutional animal care and use committees of National Defense Medical College (Approval number: 19027 and 23065). Pregnant Wistar rats and wild-type mice (C57BL/6JJcl) were purchased from CLEA Japan, Inc. (Tokyo, Japan). The day a vaginal plug was observed was regarded as embryonic day0 (e0). For sample collection from the rat neonates from postnatal day0 (P0) to P7 of birth, fetuses were vaginally delivered and were raised by the mother rat. For sample collection from the rat neonates at 6 h or 10 h of birth, full-term neonates (e21) were delivered by cesarean section and were kept in a cage warmed with a heater pad at 37°C. *Lys-EGFP* mice (20) were kindly provided by Prof. Masaru Ishii (Osaka University, Japan) and were maintained under the specific-pathogen free (SPF) conditions. The estrous cycle of mice was synchronized by the method reported by Hasegawa *et al* (57). Briefly, female mice at 10 to 20 weeks of age were subcutaneously injected with 2 mg of progesterone (FujifilmWako, Tokyo, Japan) once a day in the evening (4-6 pm) for two consecutive days. Then the mice were paired with male mice (10 to 24 weeks of age) on the 2^nd^ day. We checked a vaginal plug in the morning and regarded the day as e0. On the morning of e18 (10-12 a.m.), we performed caesarean section.

### Collection of cell suspensions and flow cytometry

We collected blood from rat fetuses or neonates using heparinized hematocrit tubes after decapitation. To remove erythrocytes, we used RBC Lysis Buffer (pluriSelect Life Science, Leipzig, Germany). We added 1.2 mL of RBS Lysis Buffer to 60 μL of blood. Then the blood was vortexed for 5 s and was incubated for 15 min at 4°C. Lysed erythrocytes were removed by centrifugation at 400 x g for 10 min. To count the absolute number of each blood leukocyte population, we used CountBright Plus Absolute Counting Beads (ThermoFisher, Waltham, MA, USA). Briefly, we added 25 μL of the counting beads solution to 60 μL of blood. To collect cell suspensions from fetal or neonatal bone marrows, we removed both ends of the femurs and tibias and manually scratched out the bone marrow using forceps in a MACS buffer (PBS supplemented with 0.5% of BSA and 2 mM of EDTA). Four femurs and four tibias from two individual rats were pooled as one sample. To obtain cell suspensions from a spleen or a liver, we minced the tissue using a razor and then dissociated the tissue using a 70 μm pore size cell strainer (Corning, Corning, NY, USA). The minced tissue was pushed against the cell strainer using a sterile syringe plunger. After centrifugation, erythrocytes were removed using RBC Lysis Buffer. The total cell count was determined using a TC-20 cell counter (Bio-Rad, Hercules, CA, USA). Each cell suspension was diluted to 1.0 × 10^6^ cells/500 μL of MACS buffer. The cell suspensions were treated with anti-CD16/32 for the blocking of the Fcγ receptor and were labeled with fluorescent dye-conjugated antibodies. The antibodies used for flow cytometry are summarized in Table S1 and Table S2. Non-viable cells were labeled with 7AAD or Zombie Aqua (BioLegend, San Diego, CA, USA). We performed flow cytometry analysis using FACS Canto II or cell-sorting using FACS Aria III (BD Bioscience, Franklin Lakes, New Jersey). We confirmed the specificity of each fluorescence signal by measuring the fluorescence minus one (FMO) control. We analyzed the acquired data using FlowJo software Ver.10 (BD Bioscience).

### BrdU labelling and flow cytometry analysis of the rat neonates and the mouse fetuses

Full-term (e21) rat fetuses were delivered by caesarean section. Ten minutes or 6 h after the delivery, we intraperitoneally administered 50 μL of bromodeoxyuridine (BrdU, Selleck Chemicals, Houston, TX, USA) solution at 100 mg/kg. Equal volume of vehicle was administered to the control group. The neonates were kept in the warmed cage for 90 min until the sample collection. The rat neonates at P1 and P3 of birth (naturally delivered) were also treated with BrdU for 90 min. Bone marrow cells were collected from BrdU-treated rat neonates. Four femurs, four tibiae, four humeri, four radiuses, and four ulnae from two individual rats were pooled as one sample. Each cell suspension was diluted to 2.0 × 10^6^ cells/500 μL of MACS buffer. The cell suspensions were treated with the fluorescent dye-conjugated antibodies and were labeled with Zombie Aqua Viability Dye (BioLegend). Then the cell suspensions were fixed with 4% paraformaldehyde, permeabilized with 5% Triton X-100/PBS, and treated with DNaseI (D4513, Merk) solution (20 μg of DNaseI in 500 μL of PBS) for 1h at 37°C. The samples were treated with anti-BrdU antibody and subjected to flow cytometry.

For BrdU labelling of the mouse fetuses, wild-type pregnant mice were intraperitoneally administered 500 μL of BrdU solution at 100 mg/kg. After 90 min or 6 h of the administration, we performed caesarean section and collected the fetal livers. The left lobes of the livers were analyzed by flow cytometry as described above. For the labelling of the major hematopoietic cell lineages (T lymphocytes, B lymphocytes, monocytes/macrophages, granulocytes, NK cells, and erythrocytes), we used Pacific Blue anti-mouse Lineage Cocktail (BioLegend) following the manufacture’s protocol.

### Administration of G-CSF neutralizing antibody to pregnant mice

Anti-mouse G-CSF antibody (MAB414) and its isotype control rat IgG1 (MAB005) were purchased from R&D (Minneapolis, NE, USA). Wild type pregnant mice were intraperitoneally injected with either G-CSF Ab or control Ab at 100 μg/kg on e15 and e16. On e17, the left lobes of the fetal livers were collected by caesarean section and were subjected to flow cytometry.

### Administration of NOS inhibitors to the rat neonates

Full-term (e21) rat fetuses were delivered by caesarean section. Ten minutes after the section, we confirmed the spontaneous respiration of the pups, and then intraperitoneally administered 50 μL of NG-Nitro-L-arginine methyl ester hydrochloride (L-NAME) or N-(3-(Aminomethyl) benzyl) acetamidine (1400W) solution. L-NAME (Dojindo, Kumamoto, Japan) was administered at 10 mg/kg and 1400W (Fujifilm Wako) was administered at 2 mg/kg. Equal volume of vehicle (PBS) was administered to the control group. The neonates were kept in the warmed cage for 5 h and 50 minutes (6 h in total) until the sample collection.

### Preparation and staining of blood cell smear

A smear of each leukocyte fraction obtained from cell sorting was prepared on a slide glass using Smear Gell (GenoStaff, Tokyo, Japan). We used WGK-1 Wright-Giemsa Stain Kit (ScyTek Laboratories, Logan, UT, USA) for the Wright-Giemsa staining.

### Histology and image analysis

For immunohistochemistry, tissues were fixed in 4% paraformaldehyde and were embedded in paraffin blocks. Immunohistochemistry was performed on four μm-thick paraffin sections. Sections were deparaffinized and washed with PBS, including 0.1% of Triton-X100. Endogenous peroxidase activity was blocked by 0.3% H_2_O_2_ in methanol. Endogenous biotin was blocked using Avidin/Biotin Blocking Kit (VECTOR LABORATORIES, Burlingame, CA, USA). Antigen-retrieval for myeloperoxidase was performed using Immunosaver (Nisshin EM, Tokyo, Japan) as instructed by the manufacturer. Nonspecific antibody reaction was blocked by applying PBS containing 5% goat serum (s1000, VECTOR LABORATORIES). Each primary antibody was diluted in PBS containing 1% goat serum and was applied to a section at 4°C for 18 h. Anti-myeloperoxidase antibody (ab208670) was purchased from Abcam. (Cambridge, UK). Biotinylated rabbit IgG (BA-1000, VECTOR LABORATORIES) was used for the detection of primary antibodies, and the signal was detected using the HRP DAB Peroxidase Substrate Kit (SK-4105, VECTOR LABORATORIES). The acquisition and analysis of immunohistochemistry images were performed using the BZ-X810 microscope and the associated software (Keyence, Osaka, Japan).

### In vivo imaging Analysis of Lys-EGFP mice

We measured the fluorescence intensity of EGFP in *Lys-EGFP* mouse neonates using the IVIS LuminaXR seriesIII system (PekinElmer, Waltham, MA, USA). Considering the fact that the homogenous *Lys-EGFP* mouse-derived neutrophils exhibit 2 to 3-fold fluorescence intensity compared to the heterogenous counterparts (20), we maintained *Lys-EGFP* mice as homogenous transgenic mice and used for the experiments. For each experiment, image acquisition and processing were performed on the same settings. Whole body imaging; excitation/emission: 480/520 nm, F stop: 2, binning: medium, exposure time: 4 sec, sample height: 0.5 cm, color scale: 4.0 × 10^7^ – 1.8 × 10^8^. Organ imaging; excitation/emission: 480/520 nm, F stop: 2, binning: small, exposure time: 7 sec, sample height: 0.1 cm, color scale: 7.0 × 10^7^ – 9.0 × 10^8^. For zoomed body imaging, we used the ZFOV-24 Zoom Lens with the setting; excitation/emission: 480/520 nm, F stop: 2, binning: medium, exposure time: 4 sec, sample height: 0.5 cm for the side view and 0.7 cm for the frontal view, color scale: 5,200 – 20,000.

### RNA isolation and quantitative RT-PCR analysis

Total RNA was extracted using the RNeasy Plus Mini Kit (QIAGEN, Venlo, Netherland). Then Reverse transcription was performed using the PrimeScript RT reagent Kit (TAKARA-Bio, Shiga, Japan). Complementary DNA was subjected to quantitative RT-PCR to quantify mRNA expressions using PikoReal 96 and the PikoReal software (ThermoFisher). PCReady primers were purchased from Eurofins Genomics (Tokyo, Japan). TB Green Premix Ex Taq II (TAKARA-Bio) was used for PCReady primer-based assays. Expressions of mRNA were determined by ΔΔCt method relative to the 18S expression (5’-gtaacccgttgaaccccatt-3’, and 5’-ccatccaatcggtagtagcg-3’). Expression of rat *Csf3* mRNA (NM_017104.2) was evaluated by TaqMan Gene Expression Assay (Rn00567344_m1, ThermoFisher).

### RNA sequencing (RNA-seq) analysis

RNA-seq analysis was performed as previously described (58). Preparation of the cDNA library and RNA-seq were performed at Genome Lead Corp. (Kagawa, Japan). The mRNA was purified using KAPA mRNA Capture Kit (KK8440, Kapa Biosystems, Wilmington, MA, USA). The cDNA library was prepared using MGIEasy Fast RNA Library Prep Set (940-000889-00, MGI, Shenzhen Shi, China). Directional RNA-seq (150bp x 2) was run on DNBSEQ-G400RS (MGI). We performed RNA-seq data analysis on RaNA-seq (https://ranaseq.eu/) (59). We used the DESeq2 R package and performed the Wald test to identify the differentially expressed genes between 10 min and 3 h. The cutoff of *P* value was set at 10 × 10^−12^.

### Western blot analysis

Protein expression was analyzed as previously reported (58). Harvested tissues were snap frozen in liquid nitrogen and were crushed using SK mill (Tokken, Chiba, Japan). Proteins were extracted with RIPA lysis buffer supplemented with the protease inhibitor cocktail (Takara-Bio, Shiga, Japan). The lysate was sonicated using an ultrasonic homogenizer (Q55, QSONICA, Newtown, CT, USA), then was centrifuged at 13,500 rpm for 10 min. The supernatant was collected, and the protein concentration was determined using the DC protein assay (Bio-Rad). We performed SDS-PAGE and western blotting using the mini-protean Tetra cell (Bio-Rad). Blotted PVDF membranes (Merk, Rahway, NJ, USA) were immersed in ice-cold acetone for fixation (60). For the membrane blocking and the antibody dilution, we used TBS-T containing 3% of bovine serum albumin. Anti-iNOS antibody (18985-1-AP) was purchased from Proteintech. Anti-GAPDH antibody (ab9485) was purchased from Abcam. HRP-conjugated secondary antibodies (anti-rabbit IgG, A24537, and anti-mouse IgG, A24524; ThermoFisher) were used for signal detection. Clarity Western ECL Substrate (Bio-Rad) was used for chemiluminescence imaging using ChemiDoc Touch (Bio-Rad). Band density was quantified using Image Lab software version 5.2 (Bio-Rad).

### Statistical analysis

Data are presented as mean + standard error of the mean (SEM) of independent experiments unless otherwise described. Data were analyzed using the unpaired Student’s *t*-test or one-way analysis of variance (ANOVA) followed by Tukey’s multiple comparison test, unless otherwise described, on GraphPad Prism version 8.4.3 (GraphPad, San Diego, CA, USA).

## Acknowledgments and funding sources

We thank Yumi Mitsui for the maintenance of the *Lys-EGFP* mice and for the technical assistance in cryosection analysis. We thank Kiyono Aoki for the contributions to the laboratory operations. This work was supported by a grant-in-aid from Kawano Masanori Memorial Public Interest Incorporated Foundation for Promotion of Pediatrics to Ryo Ishiwata and from Takeda Science Foundation to Ryo Ishiwata.

## Disclosures

No conflicts of interest, financial or otherwise, are declared by the authors.

## Author Contributions

RI and YM conceptualized and designed the study. RI designed the experiments, acquired the data, and analyzed the data. RI drafted the manuscript. YM reviewed and edited the manuscript.

---

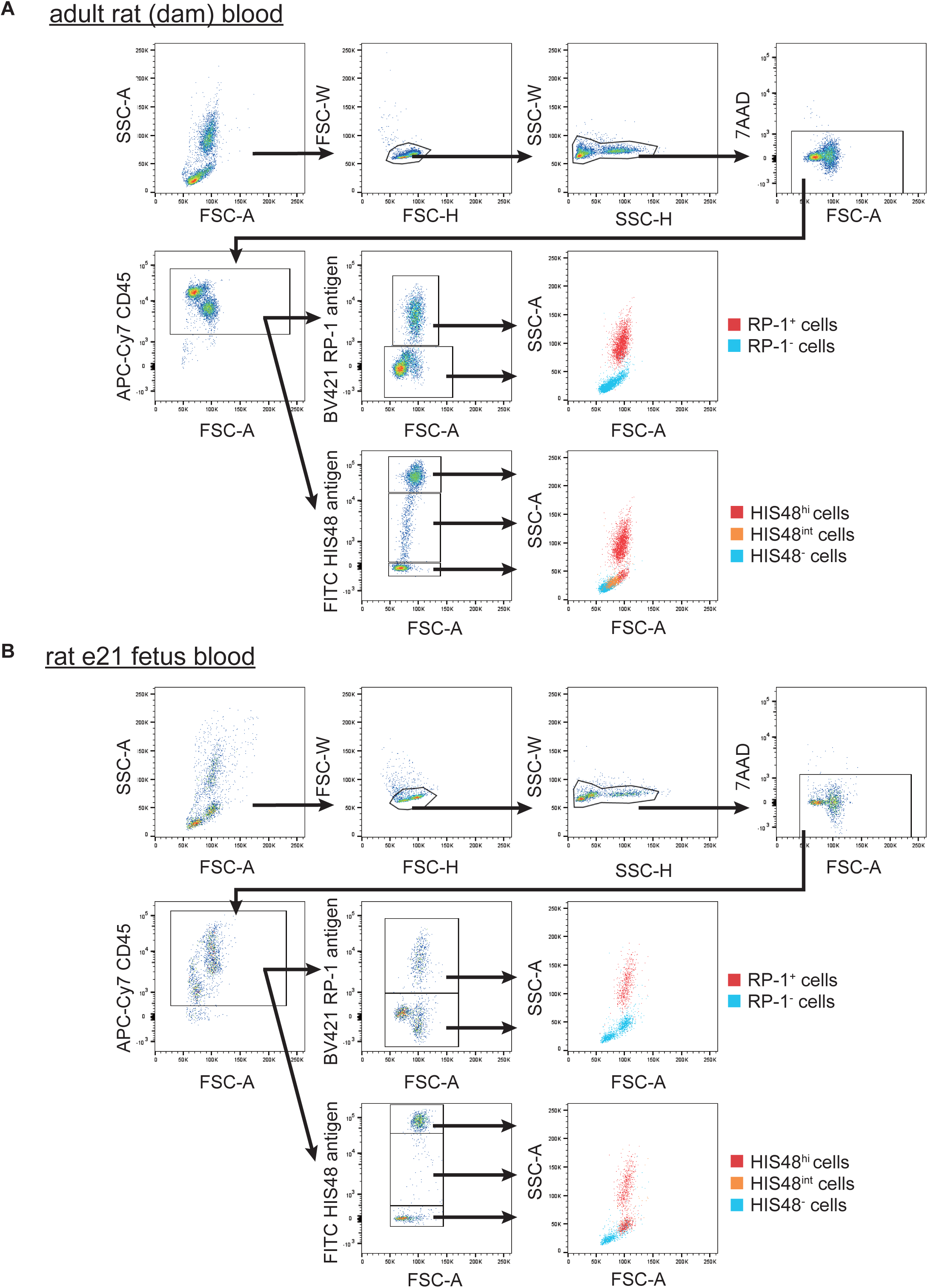

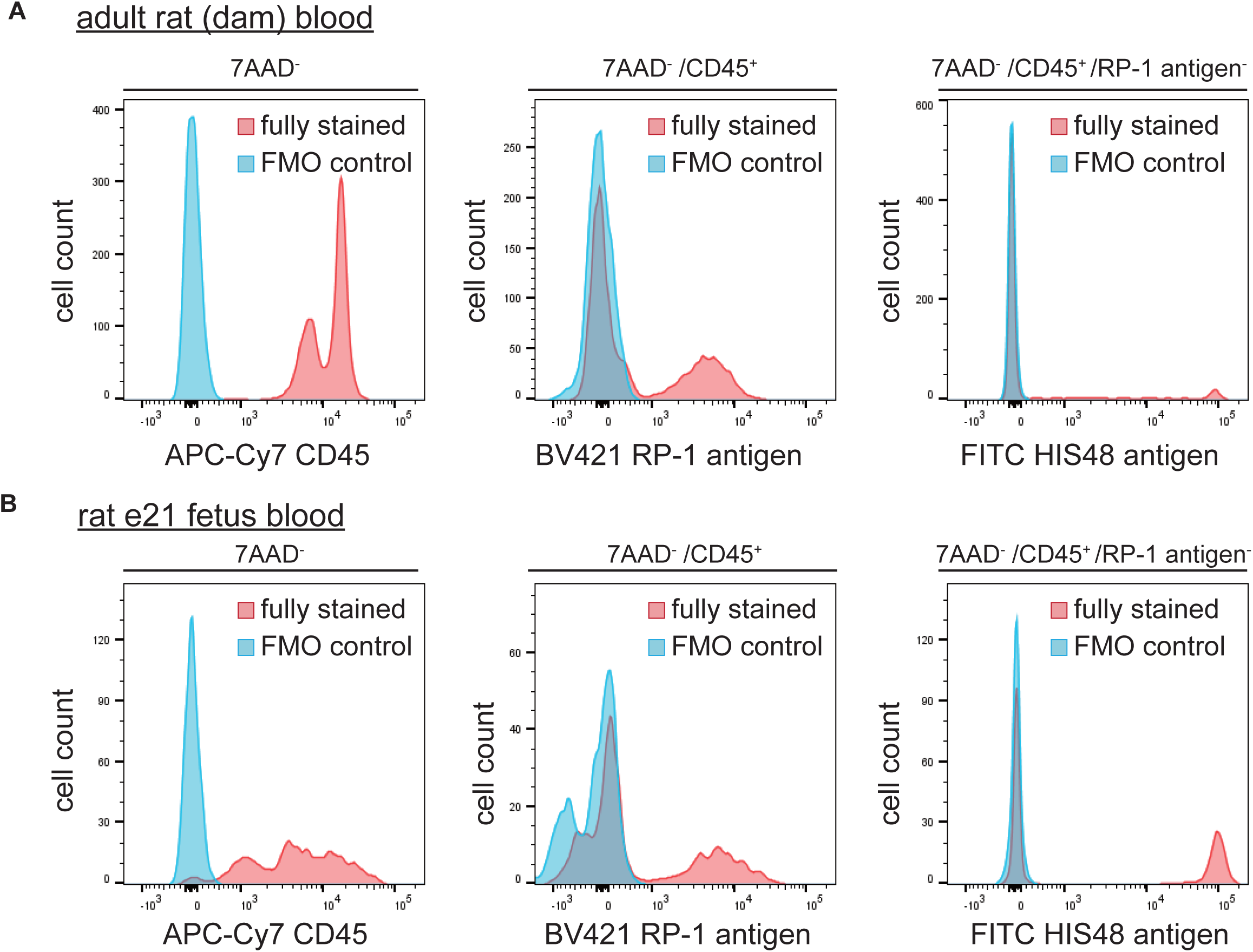

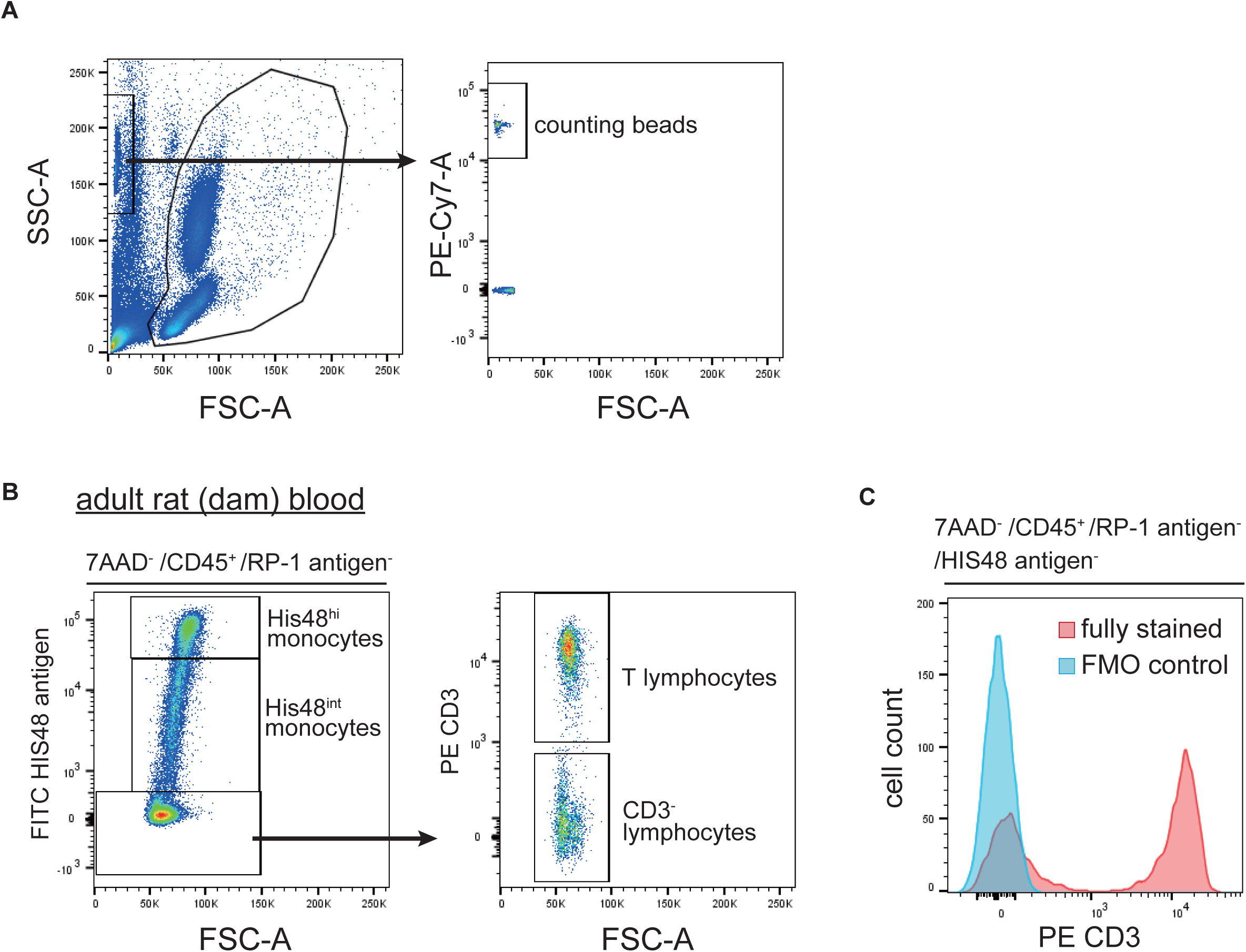

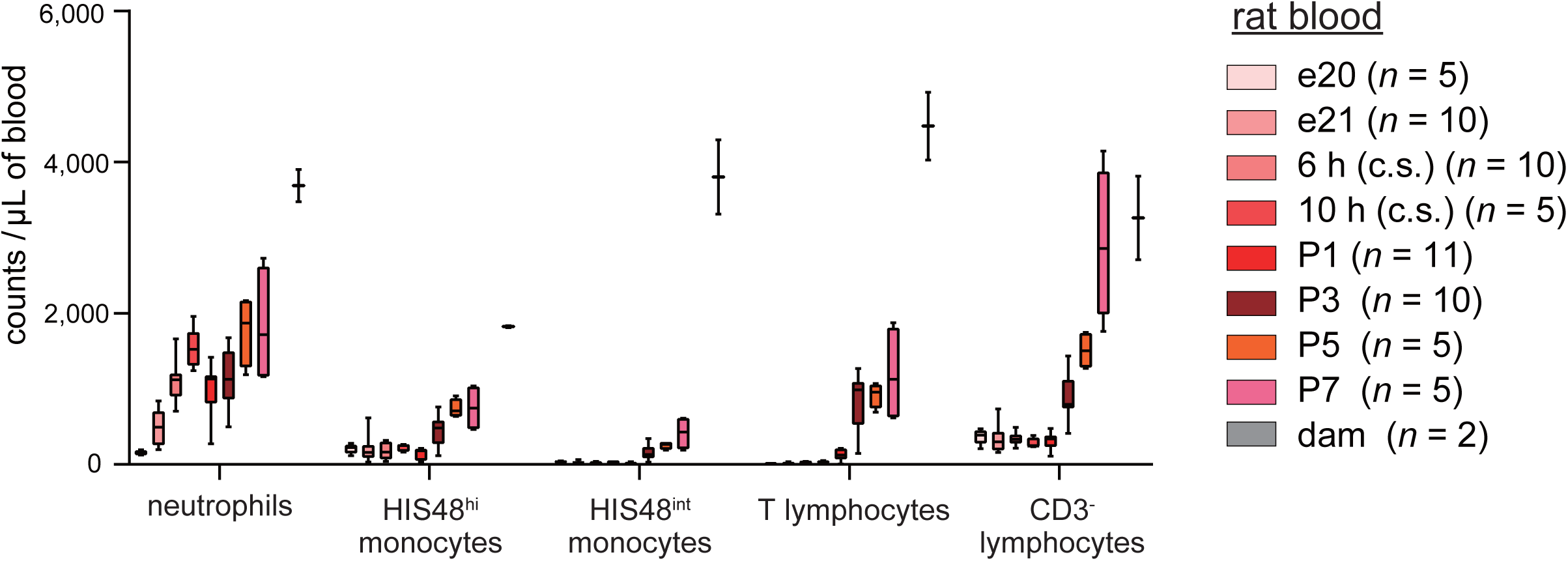

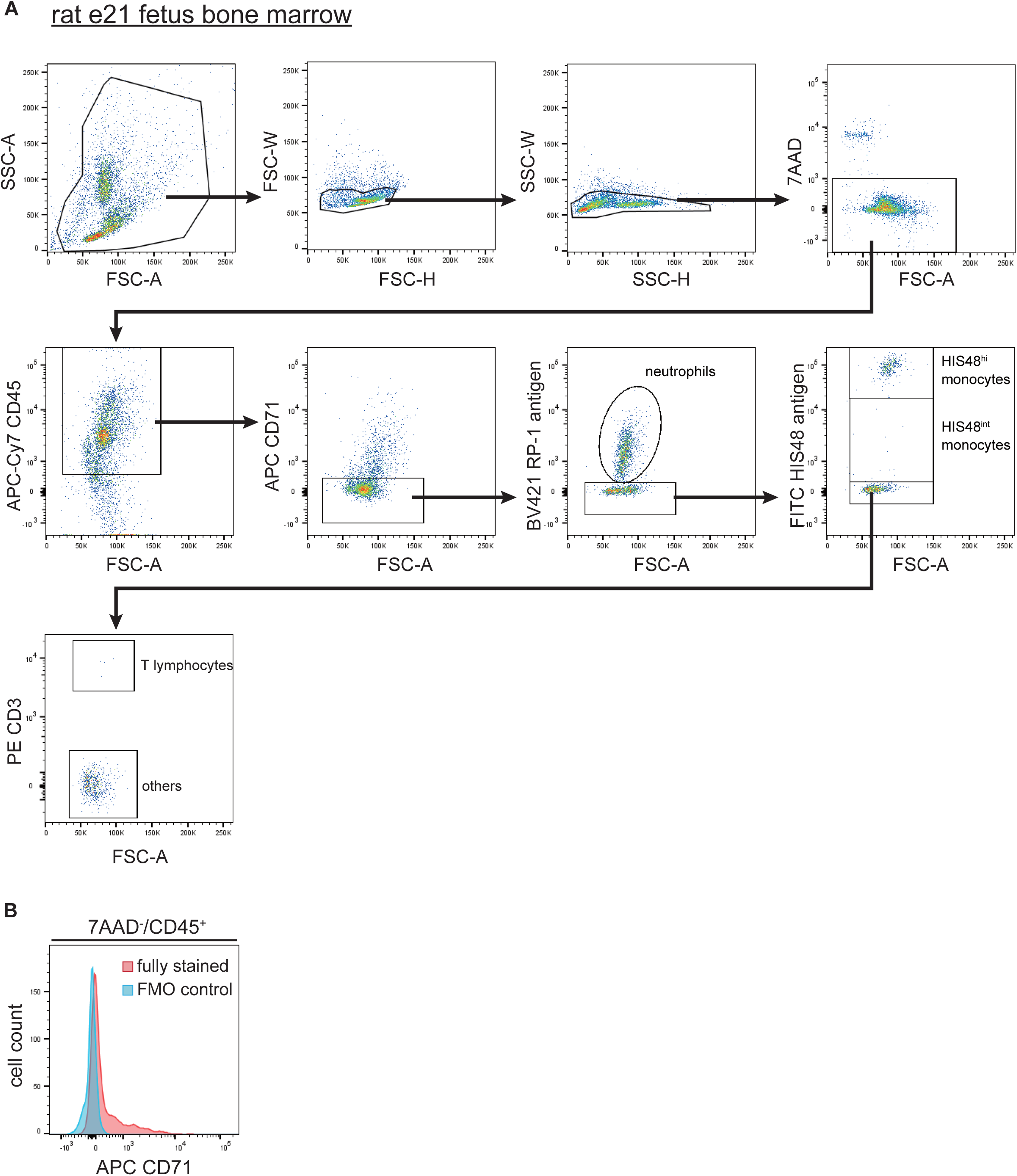

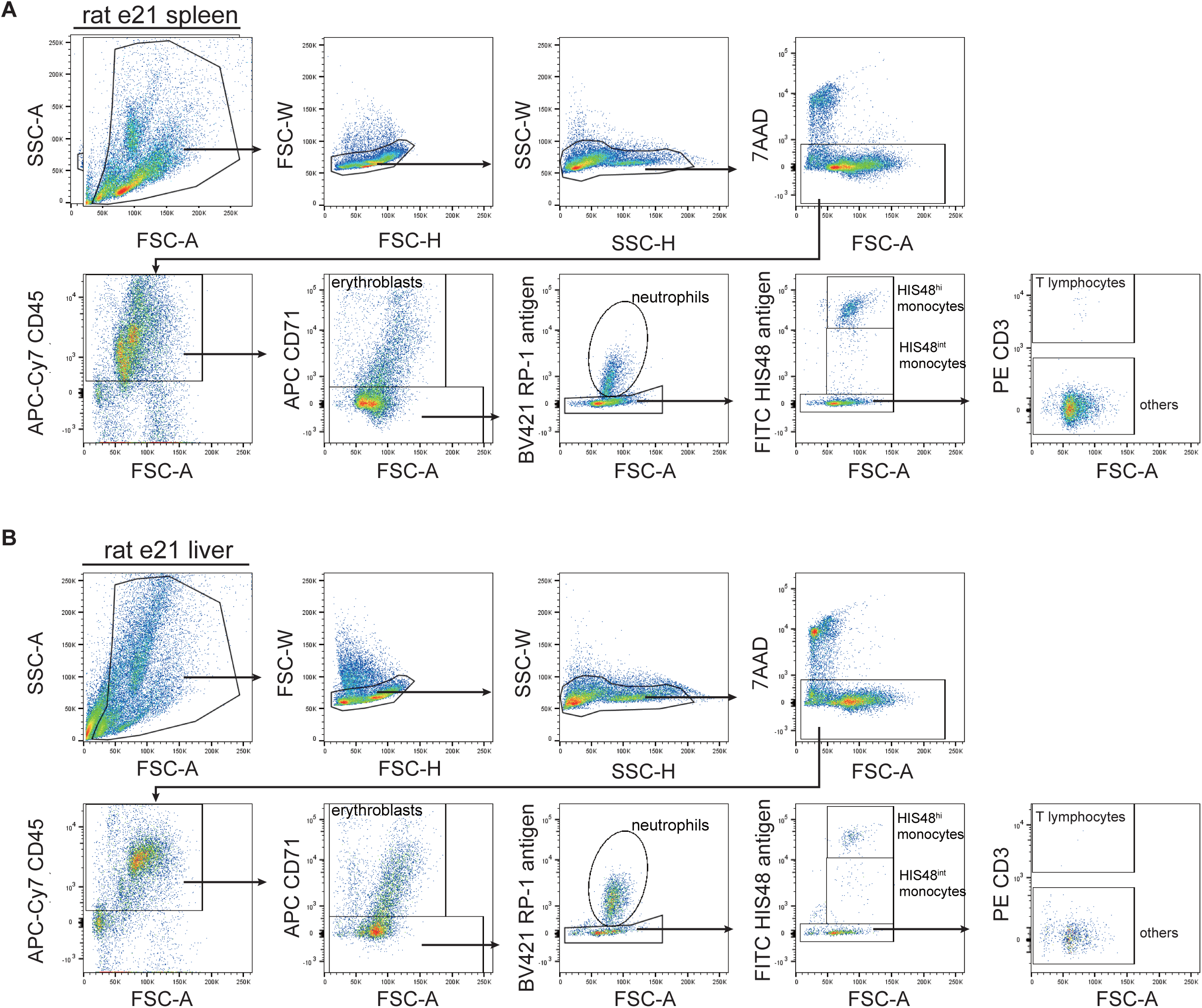

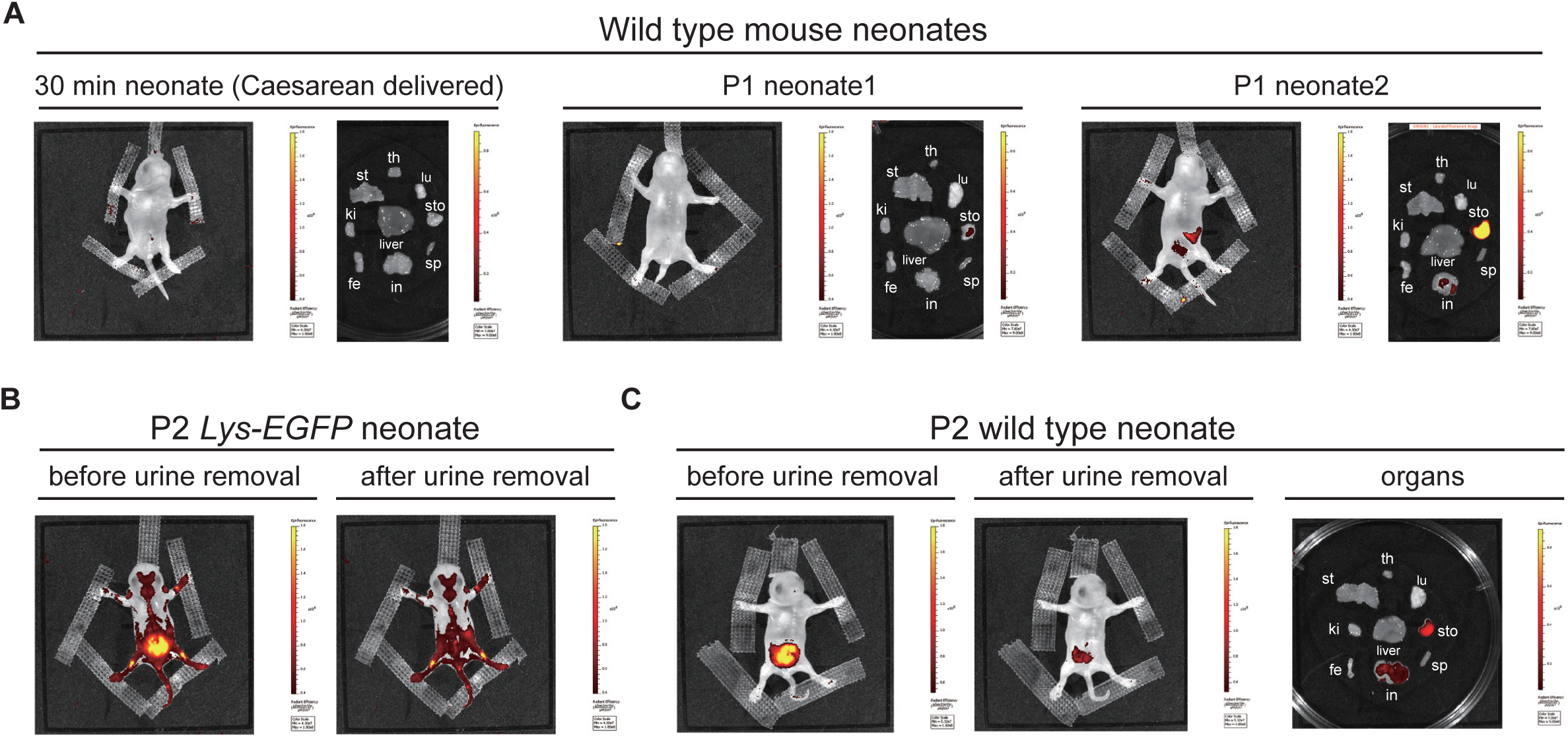

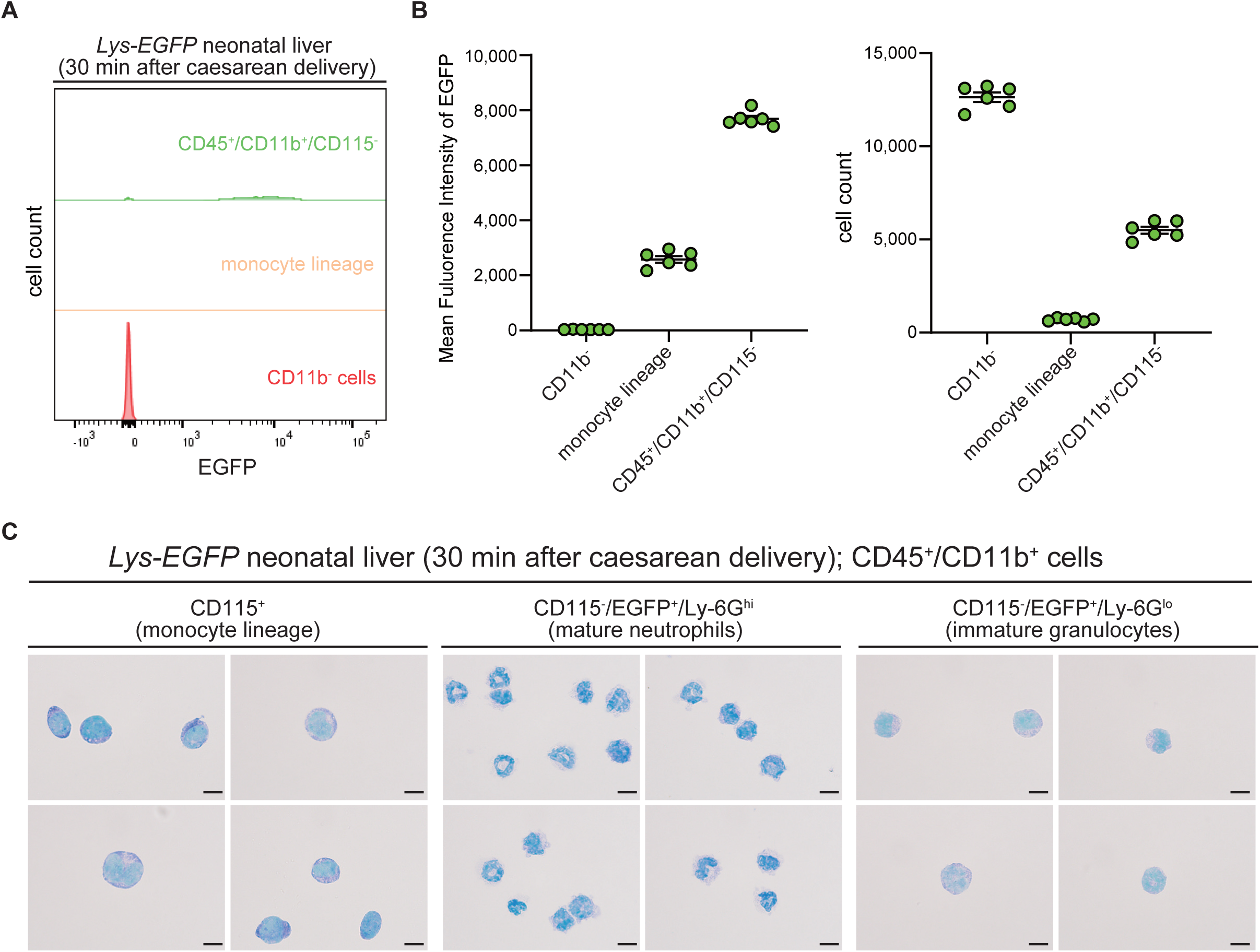

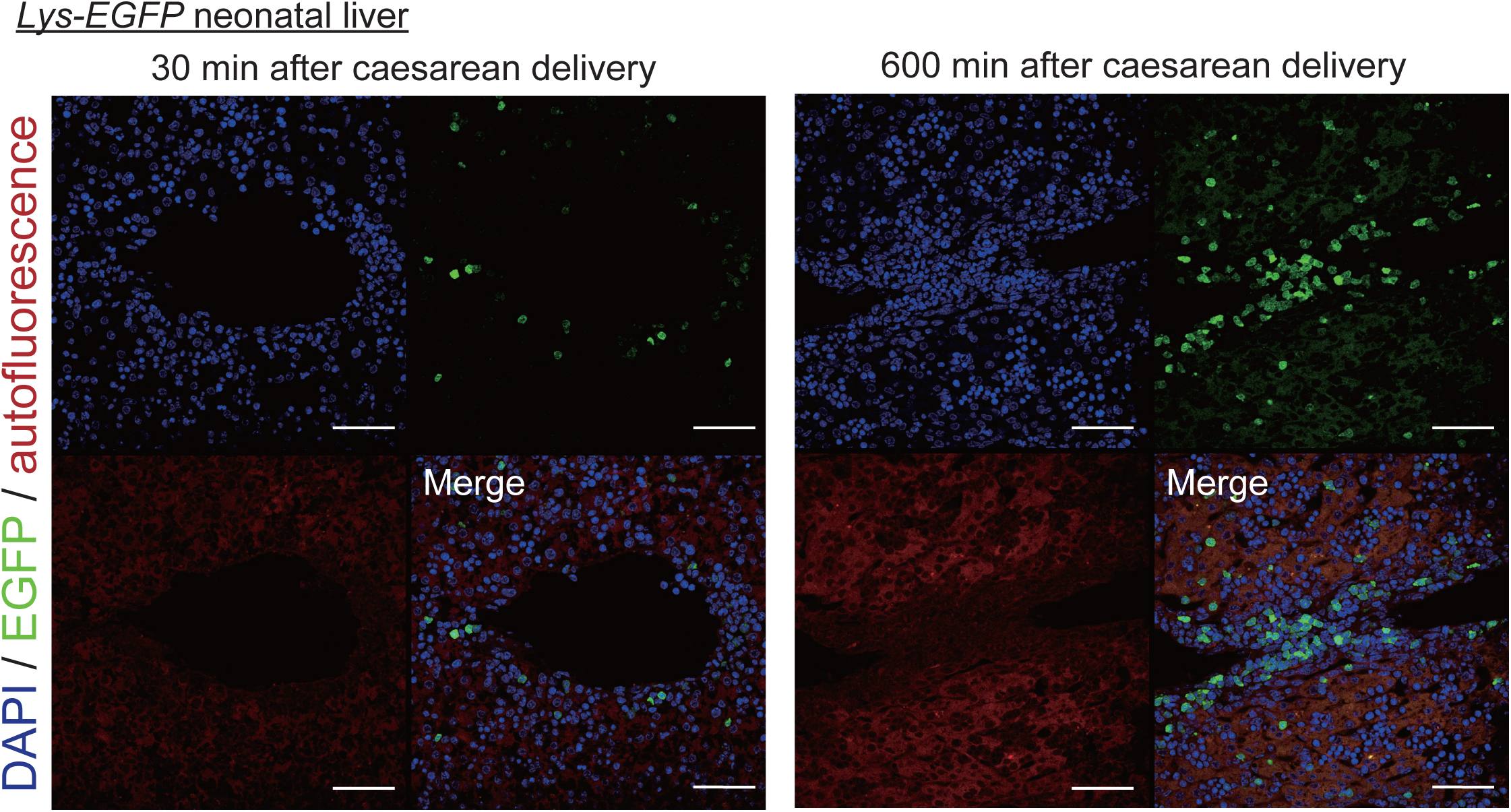

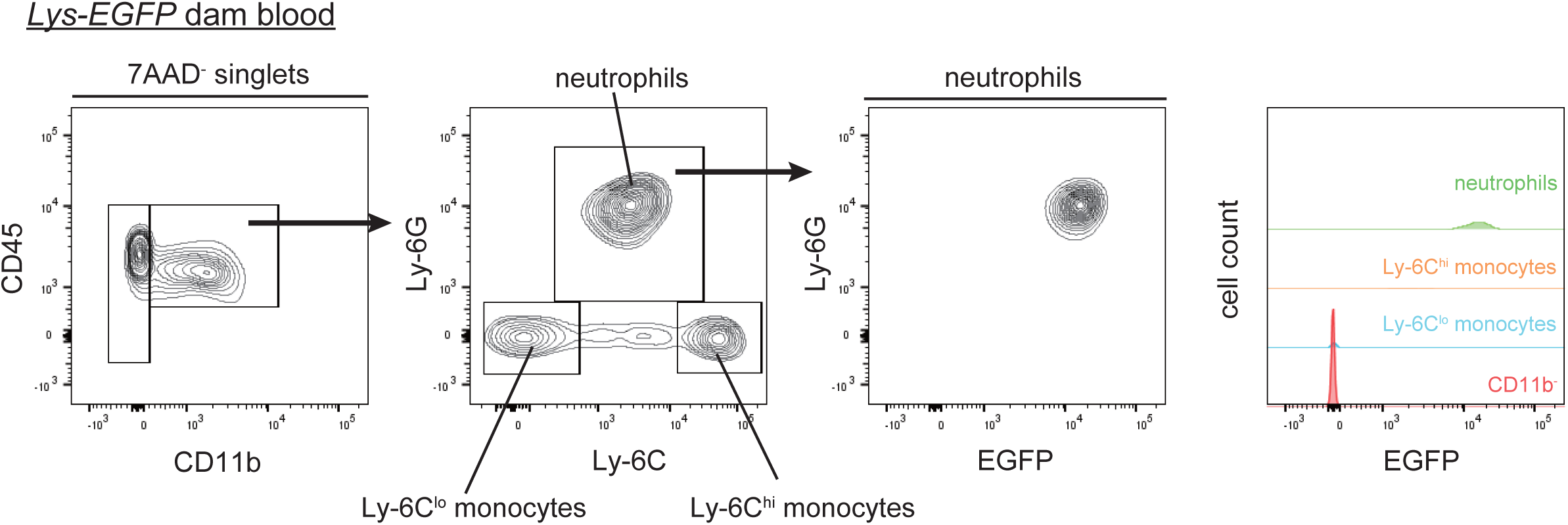

## Notes

### Competing Interest Statement

The authors have declared no competing interest.

