## Supplemental information for "Fetal liver neutrophils are responsible for the postnatal neutrophil surge"

Corresponding Author

Tel; (81)4-2995-1211

**Supplemental materials and methods**

***Histological analysis of Lys-EGFP neonatal liver***

For histological analysis of the liver tissues of the *Lys-EGFP* mouse neonates, we prepared cryosections and observed them by confocal microscopy. Briefly, the tissues were immersed in 4% paraformaldehyde overnight, then in PBS containing 30% sucrose until the tissues sink. The tissues were embedded in O.C.T compound (Sakura Finetek, Tokyo, Japan) and we prepared eight μm-thick cryosections using CM1950 cryostat (Leica, Wetzlar, Germany). The sections were mounted on CRE-12 slide glasses (MATSUNAMI, Osaka, Japan). The tissues were mounted and stained using DAPI Fluoromount-G (SouthernBiotech, Birmingham, AL, USA). Fluorescence images were obtained using TCS SP8 confocal microscope (Leica).

| Table S1. The list of antibodies used for flow cytometry of rat samples | | | | | |
| --- | --- | --- | --- | --- | --- |
| Antigen | Clone name | Dye | Manufacturer | Concentration  (mg/mL) | Dilution |
| CD32 | D34-485 | None | BD Pharmingen | 0.5 | 1-100 |
| CD45 | OX-1 | APC-Cy7 | BD Pharmingen | 0.2 | 1-150 |
| Granulocytes | RP-1 | BV421 | BD Pharmingen | 0.2 | 1-75 |
| Granulocytes | HIS48 | FITC | BD Pharmingen | 0.5 | 1-400 |
| CD3 | 1F4 | PE | BioLegend | 0.2 | 1-200 |
| CD71 | OX-26 | APC | BioLegend | 0.2 | 1-100 |
| BrdU | 3D4 | PE-Cy7 | BioLegend | 0.05 | 1-150 |

| Table S2. The list of antibodies used for flow cytometry of mouse samples | | | | | |
| --- | --- | --- | --- | --- | --- |
| Antigen | Clone name | Dye | Manufacturer | Concentration  (mg/mL) | Dilution |
| CD16/32 | 93 | None | BioLegend | 0.5 | 1-100 |
| CD45.2 | 104 | BV421 | BioLegend | 0.2 | 1-100 |
| CD45.2 | 104 | PerCP Cy5.5 | BioLegend | 0.2 | 1-100 |
| CD11b | M1/70 | BV510 | BioLegend | 0.2 | 1-100 |
| CD11b | M1/70 | PerCP Cy5.5 | BioLegend | 0.2 | 1-100 |
| Erythroid cells | TER-119 | PE | BioLegend | 0.2 | 1-200 |
| CD71 | RI7217 | APC | BioLegend | 0.2 | 1-250 |
| CD115 | AFS98 | PE-Cy7 | BioLegend | 0.2 | 1-400 |
| Ly-6G | 1A8 | APC-Cy7 | BioLegend | 0.2 | 1-400 |
| Ly-6C | HK1.4 | PE | BioLegend | 0.2 | 1-2,000 |
| Sca-1 | D7 | PE | BioLegend | 0.2 | 1-200 |
| c-kit | 2B8 | PE | BioLegend | 0.2 | 1-200 |
| c-kit | 2B8 | PE-Cy7 | BioLegend | 0.2 | 1-200 |
| CD34 | HM34 | APC | BioLegend | 0.2 | 1-100 |
| CD16/32 (FcγR) | 93 | APC-Cy7 | BioLegend | 0.2 | 1-100 |
| BrdU | 3D4 | FITC | BioLegend | 0.05 | 1-150 |

**Supplemental Figure Legends**

**Figure S1.** **Anti-rat granulocytes antibody (RP-1) labeled blood neutrophils of an adult rat and fetal rat.** Peripheral blood was collected (A) from an adult rat (dam) or from (B) a rat fetus at embryonic day 21 (e21) and was analyzed by flow cytometry**.** Singlet cells were gated on FSC-W and FSC-H plots and then on SSC-W and SSC-H plots. Dead cells (7AAD^+^ cells) were excluded. CD45^+^ leukocytes were gated by the reactivity to the two different anti-rat granulocytes antibodies, RP-1 and HIS48. RP-1^+^ and RP-1^-^ cells, or HIS48^hi^, HIS48^int^, and HIS48^-^ cells, were plotted on FSC-A and SSC-A plots.

**Figure S2. Fluorescence minus one (FMO) control for the antibodies used in the flow cytometric analysis for rat blood.** Peripheral blood was collected (A) from an adult rat (dam) or from (B) a rat fetus at embryonic day 21 (e21) and then analyzed by flow cytometry**.** An FMO control for anti-CD45, RP-1, or HIS48 was plotted against a fully stained sample.

**Figure S3. Representative flow cytometric analysis of counting beads and blood T lymphocytes.** Peripheral blood was collected from an adult rat (dam) and was analyzed by flow cytometry. (A) A representative image of the flow cytometric analysis of the counting beads. The PE-Cy7 filter was used for the detection of the fluorescent counting beads. (B) Within 7AAD^-^/CD45^+^/RP-1 ^-^ cells, the anti-CD3 antibody discriminated T lymphocytes from other lymphocytes. (C) An FMO control for anti-CD3 was plotted against a fully stained sample.

**Figure S4. The changes in the absolute leukocytes count per blood volume of rat from e20 to P7.** Changes in absolute counts per blood volume for neutrophils, HIS48^hi^ monocytes, HIS48^int^ monocytes, T lymphocytes, and CD3^-^ lymphocytes from e20 to P7 and the dams.

**Figure S5. Representative flow cytometry plots for rat fetal bone marrow cells.** (A, B) Bone marrow cells were obtained from femurs and tibiae of rat fetuses at e21 and were analyzed by flow cytometry. (A) Singlet cells were gated on FSC-W and FSC-H plots and then on SSC-W and SSC-H plots. Dead cells (7AAD^+^ cells) were excluded. Leukocytes were gated as CD45^+^ cells. Erythroblasts (CD71^+^ cells) were excluded. Neutrophils were gated as RP-1^+^ cells and monocytes as RP-1^-^/HIS48^+^ cells. Then T lymphocytes were gated as CD3^+^ cells. (B) An FMO control for anti-CD71 was plotted against a fully stained sample.

**Figure S6. Representative flow cytometry plots for a rat fetal spleen and liver.** (A) Splenic cells and (B) hepatic cells were obtained from a rat fetus at e21 and were analyzed by flow cytometry. Singlet cells were gated on FSC-W and FSC-H plots and then on SSC-W and SSC-H plots. Dead cells (7AAD^+^ cells) were excluded. Leukocytes were gated as CD45^+^ cells. Erythroblasts (CD71^+^ cells) were excluded. Neutrophils were gated as RP-1^+^ cells and monocytes as RP-1^-^/HIS48^+^ cells. Then T lymphocytes were gated as CD3^+^ cells.

**Figure S7. Nonspecific autofluorescence signals in IVIS analysis were detected for digestive orans and urine.** (A) Representative fluorescence images of a wild-type mouse neonate at 30 min of birth (caesarean-delivered) and wild-type P1 neonates. Images of dissected organs are also shown for the wild-type neonate. (B-C) Representative fluorescence images of (B) a *Lys-EGFP* P2 neonate and (C) a wild-type P2 neonate before and after urine removal. Images of dissected organs are also shown. th: thymus, st: sternum, lu: lung, ki: kidney, sto: stomach, sp: spleen, fe: femur, in: intestine.

**Figure S8. Granulocytes predominantly exhibited EGFP signals in *Lys-EGFP* in full-term neonatal livers.** (A) Representative histogram obtained from a flow cytometry analysis of a *Lys-EGFP* neonatal liver (30 min after caesarean delivery) and (B) comparisons of EGFP signal intensity and cell count for each cell fraction. Each point represents a result of an independent pup. (C) Cell sorting analysis of liver myeloid cells from an e18 *Lys-EGFP* neonate (30 min after caesarean delivery). Within CD45^+^ / CD11b^+^ cells, CD115^+^ cells (monocyte lineage), CD115^-^ / EGFP^+^ / Ly-6G^hi^ cells (neutrophils) and CD115^-^ / EGFP^+^ / Ly-6G^lo^ cells (immature granulocytes) were sorted and stained with Wright-Giemsa stain. Scale bars; 10 μm.

**Figure S9.** **Granulocytes accumulated in perivascular region after birth.** Representative fluorescence images of *Lys-EGFP* neonatal livers at 30 min or 600 min after caesarean delivery. Cryo-sections were prepared from a left lobe of a liver and perivascular area was imaged by confocal microscopy. Scale bars; 50 μm.

**Figure S10. Flow cytometry analysis on *Lys-EGFP* adult blood.**

Representative flow cytometry plots and histogram of EGFP signals for blood from a *Lys-EGFP* adult mouse (dam).
